# VezA/vezatin facilitates proper assembly of the dynactin complex in vivo

**DOI:** 10.1101/2024.04.19.590248

**Authors:** Jun Zhang, Rongde Qiu, Sean Xie, Megan Rasmussen, Xin Xiang

## Abstract

Cytoplasmic dynein-mediated intracellular transport needs the multi-component dynactin complex for cargo binding and motor activation. However, cellular factors involved in dynactin assembly remain unexplored. Here we found in *Aspergillus nidulans* that the vezatin homolog VezA is important for dynactin assembly. VezA affects the microtubule plus-end accumulation of dynein before cargo binding and cargo adapter-mediated dynein activation, two processes that both need dynactin. The dynactin complex contains multiple components including an Arp1 (actin-related protein 1) mini-filament associated with a pointed-end sub-complex. VezA physically interacts with dynactin either directly or indirectly via the Arp1 mini-filament and its pointed-end sub-complex. Loss of VezA causes a defect in dynactin integrity, most likely by affecting the connection between the Arp1 mini-filament and its pointed-end sub-complex. Using various dynactin mutants, we further revealed that assembly of the dynactin complex must be highly coordinated. Together, these results shed important new light on dynactin assembly in vivo.

## Introduction

Cytoplasmic dynein distributes organelles/vesicles/proteins/mRNAs in eukaryotic cells, and all in vivo functions of dynein need a multi-protein complex called dynactin (Schroer, 2004). Dynactin contains a mini-filament of the actin-related protein Arp1 whose two ends are capped by capping protein and a pointed-end sub-complex containing Arp11, p62, p25 and p27 (Eckley et al., 1999; Garces et al., 1999; Karki et al., 2000; Lau et al., 2021; Schafer et al., 1994). The Arp1 mini-filament and its pointed-end proteins are important for the dynein-cargo interaction via binding to cargo adapters (Chaaban and Carter, 2022; Chowdhury et al., 2015; Drerup et al., 2017; Gama et al., 2017; Lau et al., 2021; Qiu et al., 2018; Urnavicius et al., 2015; Yeh et al., 2013; Yeh et al., 2012; Zhang et al., 2014; Zhang et al., 2011). The largest subunit of dynactin is p150^Glued^ (called p150 for simplicity) (Gill et al., 1991; Holzbaur et al., 1991), which contains an N-terminal microtubule-binding domain and also interacts with dynein intermediate chain (Karki and Holzbaur, 1995; King et al., 2003; Okada et al., 2023; Singh et al., 2024; Vaughan and Vallee, 1995; Waterman-Storer et al., 1995). Another key component of dynactin is p50 (also called dynamitin) (Echeverri et al., 1996), and within dynactin, four copies of p50, two copies of p24 and two copies p150 C-termini form the dynactin shoulder (Cheong et al., 2014; Eckley et al., 1999; Maier et al., 2008; Urnavicius et al., 2015). In the presence of dynactin and cargo adapters, dynein gets activated in vitro (McKenney et al., 2014; Schlager et al., 2014). Prior to binding to dynactin and the cargo adapter, dynein most likely switches between an auto-inhibited phi conformation and an open conformation (Torisawa et al., 2014; Zhang et al., 2017a). The dynein regulator LIS1 promotes the open conformation (Karasmanis et al., 2023; Marzo et al., 2020; Qiu et al., 2019; Zhao et al., 2023), and it also promotes the formation of the activated dynein complex (Elshenawy et al., 2020; Htet et al., 2020; Singh et al., 2024). In the activated dynein complex, the dynein heavy chain tails bind to the Arp1 mini-filament as evidenced by cryo-EM studies, and some activated complexes contain two dynein dimers, both of which contact the Arp1 filaments via their tails (Grotjahn et al., 2018; Schlager et al., 2014; Urnavicius et al., 2018). In this configuration, the dynein motor domains within each dynein dimer are parallel to each other, allowing dynein to move along the microtubule (Zhang et al., 2017a). In several cell types, dynein accumulates at the microtubule plus ends, which facilitates cargo binding or cortical interaction (Han et al., 2001; Lee et al., 2003; Lenz et al., 2006; Markus et al., 2020; Sheeman et al., 2003; Splinter et al., 2012). In filamentous fungi and mammalian neurons, the plus-end accumulation of dynein needs kinesin-1 and the dynactin complex (Callejas-Negrete et al., 2015; Egan et al., 2012; Lenz et al., 2006; Penalva et al., 2017; Twelvetrees et al., 2016; Xiang et al., 2000; Yamada et al., 2010; Yao et al., 2012; Zhang et al., 2003; Zhang et al., 2008). Despite the importance of dynactin in dynein localization before cargo binding, cargo-dynein interaction and dynein activation, we know very little about how the dynactin complex is assembled in vivo.

In budding yeast where dynein is almost exclusively involved in spindle positioning (Eshel et al., 1993; Li et al., 1993), the Arp1 pointed end contains only the Arp11 homolog Arp10 but not p62, p25 or p27 (Clark and Rose, 2006; Eckley et al., 1999; Moore et al., 2008). In contrast, filamentous fungi such as *Aspergillus nidulans* and *Neurospora crassa*, where dynein is involved in both nuclear positioning and vesicle transport (Plamann et al., 1994; Seiler et al., 1999; Xiang et al., 1994), contain all four pointed-end proteins (Lee et al., 2001; Qiu et al., 2018; Zhang et al., 2008). In elongated fungal hyphae, dynein transports early endosomes and other cargos including those that hitchhike on the motile early endosomes (Egan et al., 2015; Etxebeste et al., 2013; Guimaraes et al., 2015; Higuchi et al., 2014; Lenz et al., 2006; Otamendi et al., 2019; Penalva et al., 2017; Pohlmann et al., 2015; Salogiannis et al., 2021; Salogiannis et al., 2016; Songster et al., 2023). The p25 protein at the dynactin pointed end is critical for the dynein-early endosome interaction (Zhang et al., 2011), a function conserved in mammalian cells (Yeh et al., 2012). Genetic screens have identified the endosomal dynein adapter FHF (FTS-Hook-FHIP) complex (Xu et al., 2008) including HookA in *A. nidulans* and Hok1 in *Ustilago maydis* (Bielska et al., 2014; Zhang et al., 2014). In *A. nidulans*, the cytosolic ΔC-HookA pulls down dynein-dynactin (Zhang et al., 2014), and FhipA is required for HookA to bind early endosomes (Yao et al., 2014). Function of the FHF complex in early endosome transport is conserved in mammalian cells, with Hook directly interacting with dynein-dynactin and FHIP directly interacting with Rab5 on the early endosome (Christensen et al., 2021; Guo et al., 2016; Lau et al., 2021; Olenick et al., 2019; Schroeder and Vale, 2016; Urnavicius et al., 2018). *A. nidulans* p25 is critical for the HookA-dynein-dynactin interaction (Qiu et al., 2018; Zhang et al., 2014), consistent with results from cryo-EM structural analyses on dynein-dynactin bound to mammalian Hook3 (Lau et al., 2021; Urnavicius et al., 2018).

Genetic screens in *A. nidulans* identified additional proteins important for dynein-mediated early endosome transport, including a vezatin homolog VezA (Yao et al., 2015), and Prp40A, a homolog of a splicing factor Prp40 in budding yeast and PRPF40A/PRPF40B in mammalian cells (Qiu et al., 2020). In this study, we focus on the mechanism of action of VezA/vezatin. The mammalian vezatin was initially discovered as a myosin-VIIA-binding protein involved in stabilizing adherens junctions (Kussel-Andermann et al., 2000). It was subsequently also found to be involved in neuronal functions and/or pathological conditions such as endometriosis and cancer (Bahloul et al., 2009; Danglot et al., 2012; Holdsworth-Carson et al., 2016; Koppel et al., 2019; Li et al., 2015; Pagliardini et al., 2015; Sanda et al., 2010; Sousa et al., 2004; Wang et al., 2021). Its budding yeast homolog appears to be Inp2p involved in linking myosin V to peroxisomes (Fagarasanu et al., 2006). In *A. nidulans*, the Δ*vezA* mutant exhibits an abnormal accumulation of early endosomes at the hyphal tip where microtubule plus ends are located (Yao et al., 2015). Although dynein-mediated early endosome transport still occurs in the absence of VezA, its frequency is significantly reduced (Yao et al., 2015). More recently, vezatin homologs in Drosophila and Zebrafish have been discovered as important factors for dynein-mediated axonal transport, suggesting an evolutionarily conserved role of vezatin in dynein function (Spinner et al., 2020). Here we took advantage of the *A. nidulans* genetic system to reveal the mechanism of VezA action in dynein function. Using live-cell imaging and biochemical pulldown assays in various mutant backgrounds, our results suggest that VezA is important for the proper assembly of the dynactin complex.

## Results

### VezA is important for the microtubule plus-end accumulation of dynein before cargo binding

In the Δ*vezA* mutant, early endosomes accumulate abnormally at the hyphal tip where microtubule plus ends are located (Yao et al., 2015). Although we could observe GFP-labeled dynein comets near the hyphal tip, which represent the microtubule plus-end accumulation of dynein, the dynein comets in the *vezA* mutants appeared slightly smaller than those in wild-type cells (Yao et al., 2015). Here we sought to quantify the effect of VezA on plus-end dynein accumulation, specifically in the absence of HookA (cargo adapter)-mediated dynein departure from the plus end with its cargo. To do that, we introduced the Δ*hookA* mutant allele into the Δ*vezA* mutant background and compared the signal intensity of GFP-dynein comets in the Δ*hookA*, Δ*vezA* double mutant with that in the Δ*hookA* single mutant. Although the GFP-dynein comets are present in the Δ*hookA*, Δ*vezA* double mutant, the average signal intensity is significantly lower compared to that in the Δ*hookA* single mutant (Figure 1A, 1B). This result indicates that VezA enhances the plus-end dynein accumulation, which occurs before cargo binding.

**Figure 1.**
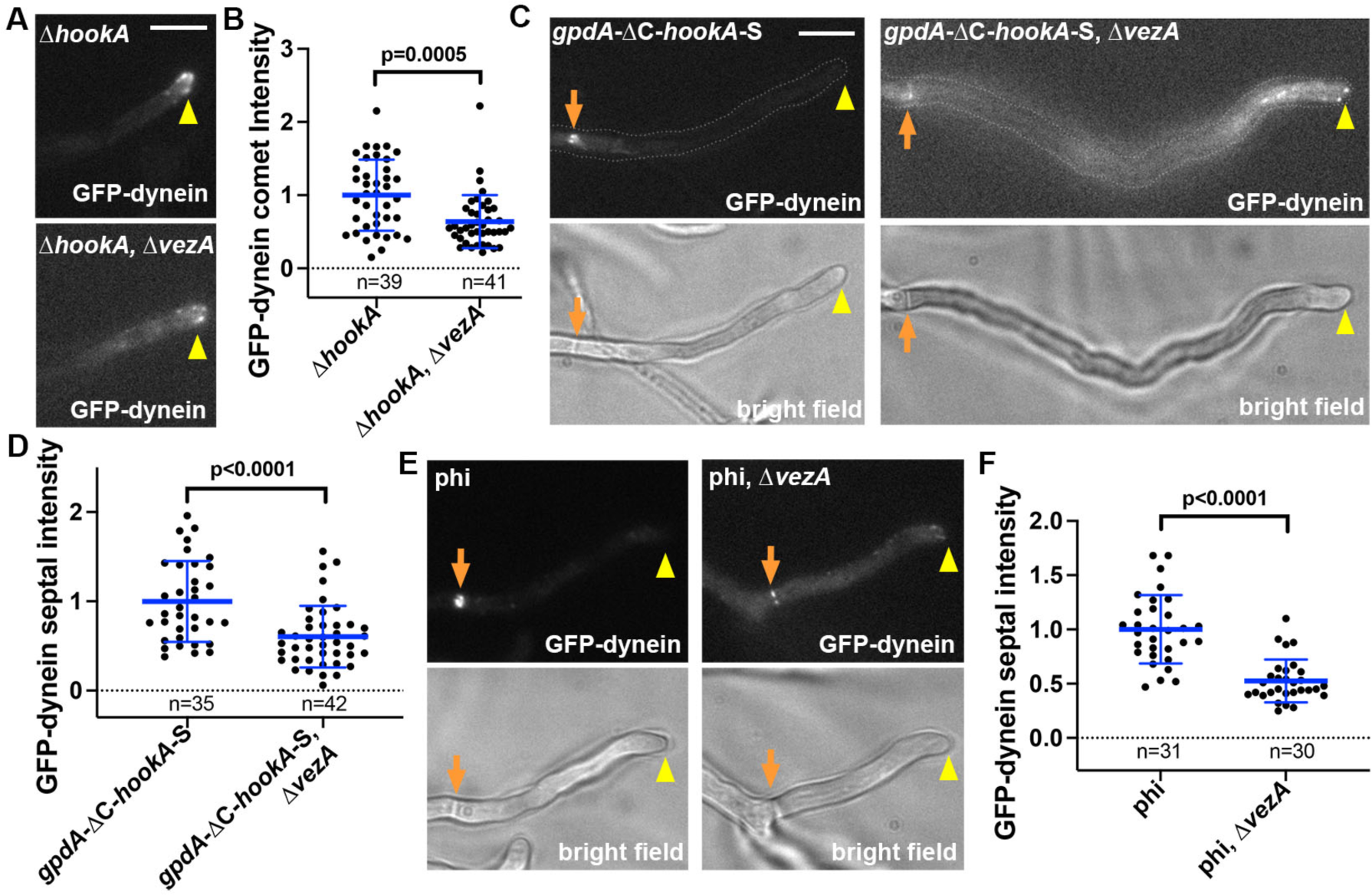
VezA affects dynein localization and cargo-adapter-mediated dynein activation. (A) Images of GFP-dynein accumulation at the microtubule plus ends as represented by comet-like structures near the hyphal tip in the Δ*hookA* and the Δ*hookA,* Δ*vezA* strains. Hyphal tip is indicated by a yellow arrowhead. Bar, 5 μm. (B) A quantitative analysis on GFP-dynein comet intensity in the Δ*hookA* and the Δ*vezA*, Δ*hookA* strains. The average value for the Δ*hookA* strain is set as 1. Scatter plots with mean and S.D. values were generated by Prism 10, and the p value was generated by Mann-Whitney test (unpaired). (C) Images of GFP-dynein in the *gpdA*-ΔC-*hookA*-S and *gpdA*-ΔC-*hookA*-S, Δ*vezA* strains. Bright-field images are shown below to indicate hyphal shape and position of septum. Hyphal tip is indicated by a yellow arrowhead and septum by a brown arrow. Bar, 5 μm. (D) A quantitative analysis on septal dynein intensity in the *gpdA*-ΔC-*hookA*-S and the *gpdA*-ΔC-*hookA*-S, Δ*vezA* strains. The average value for the *gpdA*-ΔC-*hookA*-S strain is set as 1. Scatter plots with mean and S.D. values were generated by Prism 10, and the p value was generated by Mann-Whitney test (unpaired). (E) Images of GFP-dynein in the phi mutant and phi, Δ*vezA* strains. Bright-field images are shown below to indicate hyphal shape and position of septum. Hyphal tip is indicated by a yellow arrowhead and septum by a light brown arrow. Bar, 5 μm. (F) A quantitative analysis on septal dynein intensity in the phi and phi, Δ*vezA* strains. The average value for the phi strain is set as 1. Scatter plots with mean and S.D. values were generated by Prism 10, and the p value was generated by Mann-Whitney test (unpaired).

### VezA is important for cargo adapter-mediated dynein activation

We have shown previously that overexpression of the cytosolic ΔC-HookA (HookA without the cargo-binding C-terminus) drives dynein relocation from the microtubule plus ends near the hyphal tip to septa or the spindle-pole bodies (Qiu et al., 2019), where microtubule-organizing centers (MTOCs) are located (Oakley et al., 1990; Zhang et al., 2017b). This is consistent with cargo adapter-mediated dynein activation in vitro as well as dynein activation caused by the cortical adapter Num1 in yeasts (Ananthanarayanan et al., 2013; Baumbach et al., 2017; Jha et al., 2017; Lammers and Markus, 2015; McKenney et al., 2014; Schlager et al., 2014). In *A. nidulans*, this plus-end-to-minus-end relocation causes the signal intensity of plus-end dynein comets to be diminished and the dynein signals largely accumulate at the septa (note that the accumulation at the spindle-pole bodies can only be seen at G1 (Bieger et al., 2021))(Qiu et al., 2019). To determine if VezA affects the ΔC-HookA-mediated dynein activation, we introduced into the Δ*vezA* mutant the *gpdA*-ΔC-*hookA*-S allele that causes ΔC-HookA overexpression. In the *gpdA*-ΔC-*hookA*-S, Δ*vezA* double mutant, plus-end dynein signals are more obvious than those in the *gpdA*-ΔC-*hookA*-S single mutant (Figure 1C), but the dynein signal intensity at the septal minus ends is significantly decreased compared to that in the *gpdA*-ΔC-*hookA*-S single mutant (Figure 1C, 1D). This result indicates that VezA enhances cargo adapter-mediated dynein activation in vivo.

In addition to ΔC-HookA overexpression, the phi mutations (*nudA*^R1602E,K1645E^), which overcome the autoinhibited phi conformation of dynein and keeps it in an open conformation (Zhang et al., 2017a), also drives dynein to relocate from the plus ends to the septal minus ends (Qiu et al., 2019) (Figure 1E). Importantly, deletion of *vezA* (Δ*vezA*) significantly reduced the septal accumulation of the GFP-labeled phi-mutant dynein that is constitutively open (Figure 1E, 1F). Thus, VezA must play a role beyond the phi-opening step during dynein activation. Since VezA is involved in the microtubule-plus-end accumulation of dynein before cargo binding and cargo-adapter-mediated dynein activation, two processes that both need dynactin (McKenney et al., 2014; Schlager et al., 2014; Xiang and Qiu, 2020), we further examined the relationship between VezA and the dynactin complex (Figure 2A).

**Figure 2.**
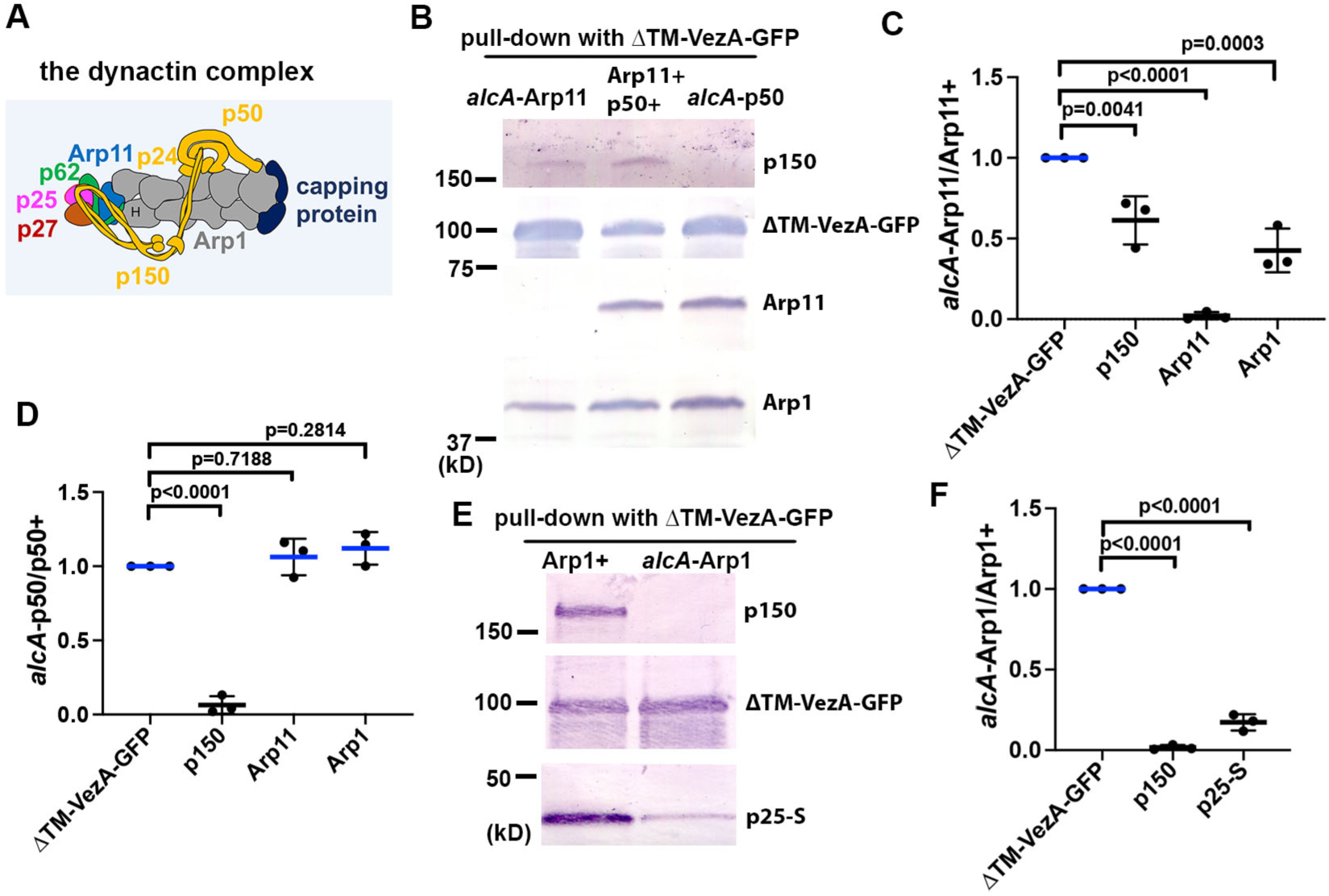
Arp1 and the pointed-end proteins are important for the interaction between VezA and dynactin. (A) A cartoon of the dynactin complex. The drawing was based on a cryo-EM structure of vertebrate dynactin (Urnavicius et al., 2015) and other publications on p150 N-terminus and its coiled-coil domains (Saito et al., 2020; Tripathy et al., 2014; Waterman-Storer et al., 1995). Except for the existence of conventional actin (labeled “H”) (Urnavicius et al., 2015) in *A. nidulans* dynactin that still needs evidence for, our previous data are consistent with the existence of all other components within *A. nidulans* dynactin (Qiu et al., 2018; Zhang et al., 2018). (B) Western blots showing dynactin components pulled down with ΔTM-VezA-GFP and the effect of shutting off the expression of Arp11 (*alcA*-Arp11) or p50 (*alcA*-p50). (C) A quantitative analysis on the effect of *alcA*-Arp11 on the amounts of other dynactin components pulled down with ΔTM-VezA-GFP. The values were generated from western blot analyses of three independent pull-down experiments (*n* = 3 for all). The ratios of the intensity value of *alcA*-Arp11 to that of wild-type Arp11 (*alcA*-Arp11/Arp11+) are shown for each component, and these ratios are relative to the ratio for ΔTM-VezA-GFP that was set as 1. Scatterplots with mean and SD values as well as p values were generated by Prism 10 (ordinary one-way ANOVA test with Dunnett’s multiple comparisons test). (D) A quantitative analysis on the effect of *alcA*-p50 on the amounts of other dynactin components pulled down with ΔTM-VezA-GFP. The values were generated from western blot analyses of three independent pull-down experiments (*n* = 3 for all). The ratios of the intensity value of *alcA*-p50 to that of wild-type p50 (*alcA*-p50/p50+) are shown for each component, and these ratios are relative to the ratio for ΔTM-VezA-GFP that was set as 1. Scatterplots with mean and SD values as well as p vales were generated by Prism 10 (ordinary one-way ANOVA test with Dunnett’s multiple comparisons test). (E) Western blots showing the effect of shutting of Arp1 expression (*alcA*-Arp1) on the amounts of p150 and p25-S pulled down with ΔTM-VezA-GFP. (F) A quantitative analysis on the effect of *alcA*-Arp1 on the amounts of p150 and p25-S pulled down with ΔTM-VezA-GFP. The values were generated from western blot analyses of three independent pull-down experiments (*n* = 3 for all). The ratios of the intensity value of *alcA*-Arp1 to that of wild-type Arp1 (*alcA*-Arp11/Arp11+) are shown for each component, and these ratios are relative to the ratio for ΔTM-VezA-GFP that was set as 1. Scatterplots with mean and SD values as well as p values were generated by Prism 10 (ordinary one-way ANOVA test with Dunnett’s multiple comparisons test).

### VezA interacts with dynactin in an Arp1- and pointed-end-dependent fashion

While the full-length VezA is hard to be extracted, overexpressed ΔTM-VezA-GFP (missing two predicted transmembrane domains (Yao et al., 2015)) pulls down dynactin in cell extract (Figure 2B). Dynein and its binding proteins are pulled down as well, as expected (Table S1; Supplemental file 1). To determine which dynactin component is involved in the interaction, we used several conditional-null mutants including *alcA*-Arp11, *alcA*-p50 and *alcA*-Arp1, in which the *alcA* promoter can be repressed by glucose (Waring et al., 1989). Loss of Arp11 (*alcA*- Arp11) caused a significant decrease in the amounts of p150 and Arp1 pulled down with ΔTM- VezA-GFP (Figure 2B, 2C), a result further supported by a mass spectrometry analysis (Table S1; Supplemental data file 1). Loss of p50 (*alcA*-p50) diminished the amount of p150 pulled down with ΔTM-VezA-GFP (Figure 2B, 2D; Table S1), but it did not significantly affect the amounts of Arp1 and Arp11 pulled down with ΔTM-VezA-GFP (Figure 2B, 2D). Consistently, a mass spectrometry analysis on ΔTM-VezA-GFP pulldown in the *alcA*-p50 extract detected no peptides of p150 but detected about normal numbers of peptides from pointed-end proteins Arp11, p62 and p25 (Table S1; Supplemental data file 1). Finally, loss of Arp1 (*alcA*-Arp1) abolished the pulldown of p150, and it also caused a significant reduction in the pulled-down amount of S-tagged p25 (p25-S) (Figure 2E, 2F). Consistently, the mass spectrometry analysis on ΔTM-VezA-GFP pulldown in the *alcA*-Arp1 extract detected no peptides of p150 and p50 but detected peptides from the pointed-end proteins Arp11, p62 and p25 (Table S1; Supplemental data file 1), suggesting that the pointed-end sub-complex is able to bind VezA. Together, our results suggest that the VezA-dynactin interaction is independent of p150 or p50 but depends on Arp1 and its pointed-end proteins.

So far, we have not been able to purify dynactin and VezA to test whether the VezA-dynactin association is direct, but we used AlphaFold2 to test whether a direct interaction is possible. VezA was predicted to form a homodimer (a prediction first made by Andrew Carter at MRC Laboratory of Molecular Biology). Thus, we used two copies of VezA in an AlphaFold2-based analysis including the pointed-end proteins (Arp11, p62, p25 and p27) and four copies of Arp1. Four models were obtained and one of them is shown (Figure S1A). In this model (as well as in two other models), the N-terminus of VezA (aa 1-20) is close to the pointed end via p62 and p25 (Figure S1B). The C-terminal α-helix (563-615) is docked in a pocket formed by two Arp1 subunits (Figure S1C). To test the importance of the N- and C-terminal regions, we made the *vezA*^Δ1-20^-GFP and *vezA*^Δ563-615^-GFP mutants. The mutant proteins are expressed although their levels are moderately lower than that of the functional full-length *vezA*^1-615^-GFP (Figure S1D, S1E). Because VezA is important for dynein-mediated early endosome transport (Yao et al., 2015), we observed early-endosome distribution in these mutants by using the Δ*vezA* and *vezA*^1-615^-GFP strains as controls. Early endosomes in the *vezA*^Δ1-20^-GFP and *vezA*^Δ563-615^-GFP mutants are abnormally accumulated at the hyphal tip just like in the Δ*vezA* mutant (Figure S1F, S1G), consistent with the functional importance of the N- and C-termini of VezA.

### VezA is important for assembly of the dynactin complex

Consistent with earlier observations in mammalian cells (Valetti et al., 1999; Vaughan et al., 1999; Vaughan et al., 2002), dynactin p150-GFP in *A. nidulans* forms plus-end comets, which depends on its microtubule-binding domain (Yao et al., 2012; Zhang et al., 2003) (Figure 3A). Similarly, Arp11-GFP (Qiu et al., 2020), p25-GFP (Zhang et al., 2011), and p62-GFP all form plus-end comets near the hyphal tip (Figure 3A; Figure S2), although their signal intensities appeared lower than that of p150-GFP, most likely because they are monomeric whereas p150 is dimeric within dynactin. In addition, p50-GFP forms bright plus-end comets (Figure 3A). In the Δ*vezA* mutant, plus-end comets of p50-GFP looked obviously dimmer, and those of Arp11-GFP could hardly be seen (Figure 3A). Similarly, those of p62-GFP and p25-GFP could hardly be seen either near the hyphal tip of the Δ*vezA* mutant (Figure S2A). Previously, we found that loss of the HookA cargo adapter enhanced the plus-end accumulation of p150, likely because it prevents dynein-dynactin from leaving the plus end (Qiu et al., 2018) (Figure 3B). Consistently, the plus-end accumulations of Arp11-GFP, p62-GFP or p25-GFP are also strong in the Δ*hookA* background (Figure 3B; Figure S2B), which facilitates the quantification of comet signal intensity in the Δ*vezA* mutant (Figure 3C; Figure S2C). We found that the plus-end accumulation of all examined dynactin components is reduced in the Δ*vezA*, Δ*hookA* double mutant compared to that in the Δ*hookA* single mutant (Figure 3C). Among the dynactin components examined, the pointed-end proteins such as Arp11-GFP, p62-GFP and p25-GFP were most severely affected by the loss of VezA (note that the mean of Arp11-GFP comet intensity in the Δ*vezA*, Δ*hookA* background is ∼36% of that in the Δ*hookA* background) (Figure 3B, 3C; Figure S2); p50-GFP was affected more severely (mean in Δ*vezA*, Δ*hookA* is ∼51% of that in Δ*hookA*) compared to p150-GFP (mean in Δ*vezA*, Δ*hookA* is ∼72% of that in Δ*hookA*) (Figure 3B, 3C).

**Figure 3.**
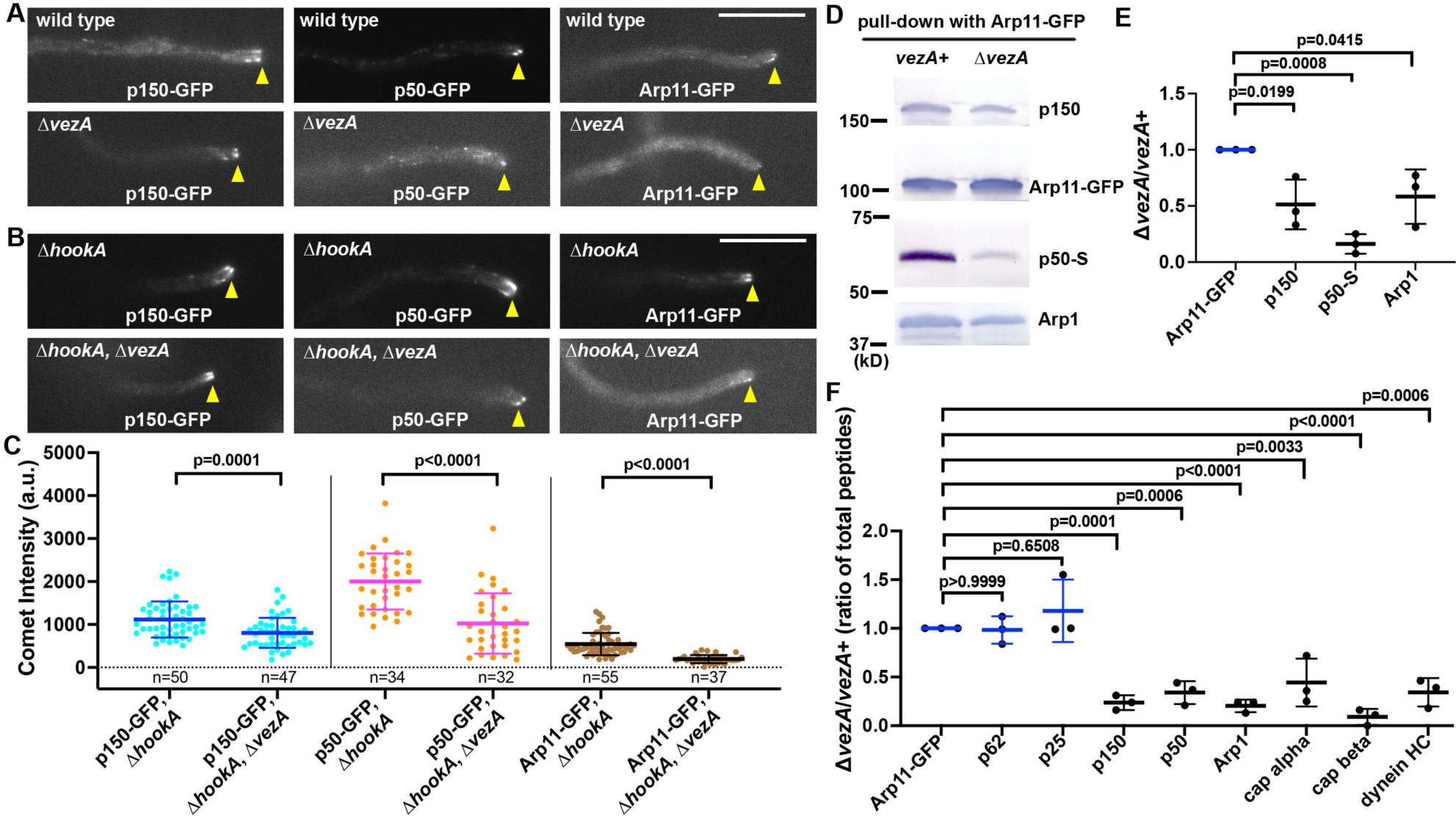
Loss of VezA causes a defect in the dynactin complex. (A) Images of p150-GFP, p50-GFP and Arp11-GFP accumulation at the microtubule plus ends as represented by comets near the hyphal tip in the wild-type and Δ*vezA* strains. Hyphal tip is indicated by a yellow arrowhead. Bar, 5 μm. (B) Images of p150-GFP, p50-GFP and Arp11-GFP in the Δ*hookA* single mutant and the Δ*hookA*, Δ*vezA* double mutant strains. Hyphal tip is indicated by a yellow arrowhead. Bar, 5 μm. (C) Quantitative analyses on plus-end comet intensity of p150-GFP, p50-GFP and Arp11-GFP in the Δ*hookA* and Δ*hookA*, Δ*vezA* strains. The average values for the intensity in the Δ*hookA* singe mutant was set as 1. Scatter plots with mean and S.D. values were generated by Prism 8, and the p values were generated by Mann-Whitney test (unpaired). (D) Western blots showing that the amounts of p150, p50-S and Arp1 pulled down with Arp11-GFP are lowered in the Δ*vezA* mutant. (E) A quantitative analysis on the effect of Δ*vezA* on the amounts of p150, p50-S and Arp1 pulled down with Arp11-GFP. The values were generated from western blot analyses of three independent pull-down experiments (*n* = 3 for all). The ratios of the intensity value of Δ*vezA* to that of wild-type *vezA* (Δ*vezA*/*vezA*+) are shown for each component, and these ratios are relative to the ratio for Arp11-GFP that was set as 1. Scatterplots with mean and SD values as well as p values were generated by Prism 10 (ordinary one-way ANOVA test with Dunnett’s multiple comparisons test). (F) A quantitative analysis on the effect of Δ*vezA* on the amounts of proteins pulled down with Arp11-GFP. The values were generated from three mass spectrometry analyses of three independent pull-down experiments (*n* = 3 for all). The ratios of the total peptide number of Δ*vezA* to that of wild-type *vezA* (Δ*vezA*/*vezA*+) for each protein are shown, and these ratios are relative to the ratio for Arp11-GFP that was set as 1. Scatterplots with mean and SD values as well as p values were generated by Prism 10 (ordinary one-way ANOVA test with Dunnett’s multiple comparisons test).

The significant reduction in the microtubule plus-end accumulation of pointed-end proteins suggested a defect in their association with p150, which contains the microtubule-binding domain critical for its microtubule plus-end accumulation (Kim et al., 2007; Lloyd et al., 2012; Moughamian and Holzbaur, 2012; Vaughan et al., 2002; Yao et al., 2012). To examine this notion further, we used Arp11-GFP for protein pulldown assays using extracts from the wild-type and Δ*vezA* strains. Loss of VezA caused a significant reduction in the amounts of p150, p50-S (S-tagged p50) and Arp1 pulled down with Arp11-GFP (Figure 3D, 3E). Furthermore, our mass spectrometry analysis showed that the amounts of p150, p50, Arp1 and capping protein pulled down with Arp11-GFP were all decreased in Δ*vezA* samples (Figure 3F; Table S2; Supplemental data file 2), consistent with a defect in dynactin assembly. Arp11-GFP also pulled down much lower amount of dynein in the Δ*vezA* extract (Figure 3F; Table S2), which is expected since p150 and Arp1 are critical for binding dynein (Chowdhury et al., 2015; Karki and Holzbaur, 1995; Singh et al., 2024; Urnavicius et al., 2015; Vaughan and Vallee, 1995). However, the amounts of pointed-end proteins p62 and p25 pulled down with Arp11-GFP were not significantly affected (Figure 3F; Table S2), suggesting that the pointed-end sub-complex is stable upon loss of VezA.

### Loss of a dynactin component can affect the integrity of other parts of the complex

In the Δ*vezA* mutant, some functional dynactin complexes must still exist, because dynein-mediated transport of early endosomes still occurs albeit at a significantly lowered frequency (Yao et al., 2015) and the Δ*vezA* colony is healthier than dynactin mutants (Yao et al., 2015; Zhang et al., 2008). Consistently, dynein-mediated nuclear distribution is only partially defective in the Δ*vezA* mutant (Figure S3), as compared to the *nudA*1 dynein mutant examined under the same conditions (Zhang et al., 2023). Thus, VezA is not essential for dynactin assembly but facilitates this process to ensure optimal dynein function.

Our finding on the involvement of VezA in dynactin assembly stimulated us to further explore how other dynactin components affect the integrity of the complex in vivo. Our current result (Figure 3F) suggested that the pointed-end sub-complex is stable on its own. This notion is consistent with previous results that the sub-complex is stable upon separation from the purified dynactin complex (Eckley et al., 1999) and that the pointed-end proteins can be purified as a complex from cultured cells expressing the recombinant proteins (Gama et al., 2017; Lau et al., 2021). Thus, we continued to use GFP-fusion of a pointed-end protein for pulldown assays as its level is unlikely to be affected by the loss of other parts of dynactin. We found that loss of p50 (*alcA*-p50) significantly reduced the amounts of p150 and Arp1 pulled-down with p25-GFP although the amount of Arp11 pulled-down was not affected significantly (Figure 4A, 4B). This effect on p150 may be caused by a role of p50 in maintaining p150 stability or in strengthening the p150-Arp1 interaction. The effect on Arp1 suggests that loss of p50 affects the assembly of the Arp1 mini-filament and/or its interaction with the pointed-end sub-complex in vivo. We further examined the effect of the *alcA*-p150 allele and found that loss of p150 significantly decreased the amounts of p50 and Arp1 pulled-down with p25-GFP (Figure 4C, 4D). Thus, p50 stability depends on p150, and the shoulder complex is important for the proper assembly of the Arp1 mini-filament and/or its interaction with the pointed-end sub-complex in vivo.

**Figure 4.**
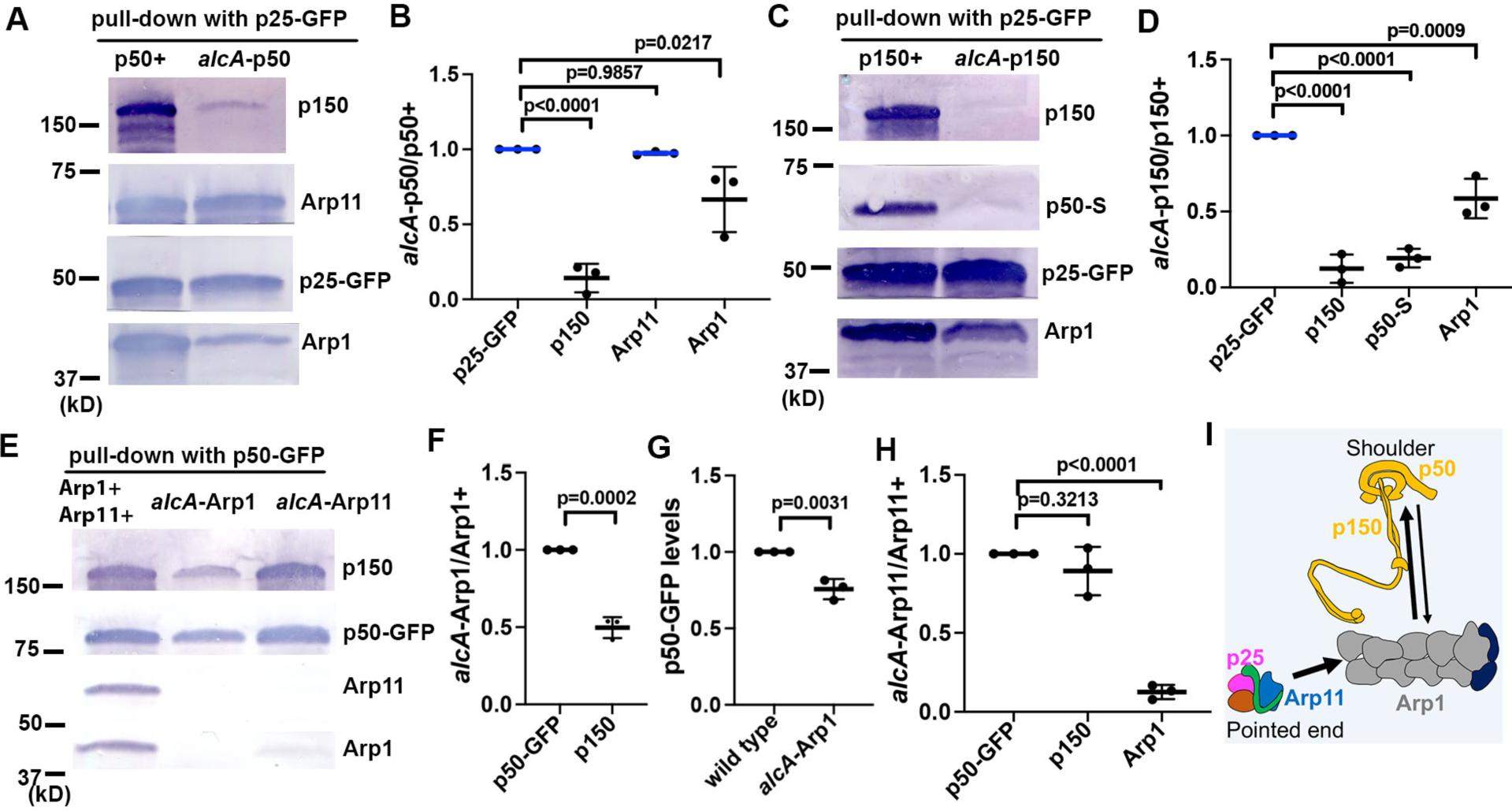
Various dynactin parts affect the integrity of each other. (A) Western blots showing that upon loss of p50 (*alcA*-p50), the amounts of p150 and Arp1 pulled down with p25-GFP are lowered but that of Arp11 is not. (B) A quantitative analysis on the effect of *alcA*-p50 on the amounts of p150, Arp1 and Arp11 pulled down with p25-GFP. The values were generated from western analyses of three independent pull-down experiments (*n* = 3 for all). The ratios of the intensity value of *alcA*-p50 to that of wild-type p50 (*alcA*-p50/p50+) are shown for each component, and these ratios are relative to the ratio for p25-GFP that was set as 1. Scatterplots with mean and SD values as well as p values were generated by Prism 10 (ordinary one-way ANOVA test with Dunnett’s multiple comparisons test). (C) Western blots showing that upon loss of p150 (*alcA*-p150), the amounts of p50-S and Arp1 pulled down with p25-GFP are lowered. (D) A quantitative analysis on the effect of *alcA*-p150 on the amounts of p150, p50-S and Arp1 pulled down with p25-GFP. The values were generated from western blot analyses of three independent pull-down experiments (*n* = 3 for all). The ratios of the intensity value of *alcA*-p150 to that of wild-type p150 (*alcA*-p150/p150+) are shown for each component, and these ratios are relative to the ratio for p25-GFP that was set as 1. Scatterplots with mean and SD values as well as p values were generated by Prism 10 (ordinary one-way ANOVA test with Dunnett’s multiple comparisons test). (E) Western blots showing that the amount of p150 pulled down with p50-GFP as well as the level of p50-GFP itself are lowered upon loss of Arp1 (*alcA*-Arp1), and that the amount of Arp1 pulled down with p50-GFP is lowered upon loss of Arp11 (*alcA*-Arp11). (F) A quantitative analysis on the effect of *alcA*-Arp1 on the amounts of p150 pulled down with p50-GFP. The values were generated from western analyses of three independent pull-down experiments (*n* = 3 for all). The ratios of the intensity value of *alcA*-Arp1 to that of wild-type Arp1 (*alcA*-Arp1/Arp1+) are shown, and the p150 ratio is relative to the p50-GFP ratio that was set as 1. Scatterplots with mean and SD values as well as p values were generated by Prism 10 (Student’s *t* test, two tailed, unpaired). (G) A quantitative analysis on the effect of *alcA*-Arp1 on the level of p50-GFP. The values were generated from western blot analyses of three independent pull-down experiments (*n* = 3 for all). Scatterplots with mean and SD values as well as p values were generated by Prism 10 (Student’s *t* test, two tailed, unpaired). (H) A quantitative analysis on the effect of *alcA*-Arp11 on the amounts of Arp1 pulled down with p50-GFP. The values were generated from western blot analyses of three independent pull-down experiments (*n* = 3 for all). The ratios of the intensity value of *alcA*-Arp1 to that of wild-type Arp1 (*alcA*-Arp1/Arp1+) are shown, and the ratios are relative to the p50-GFP ratio that was set as 1. Scatterplots with mean and SD values as well as p values were generated by Prism 10 (ordinary one-way ANOVA test with Dunnett’s multiple comparisons test). (I) A cartoon illustrating that the shoulder and Arp1 mini-filament affect each other (arrows) to control the integrity of both parts, and that the pointed-end sub-complex affects the integrity of the Arp1 mini-filament (arrow).

Previous data suggested that Arp1 is required for p150 stability (Haghnia et al., 2007; Minke et al., 1999; Zhang et al., 2008) and Arp11 is required for integrity of the dynactin complex (Clark and Rose, 2006; Yeh et al., 2012; Zhang et al., 2008). Here we performed p50-GFP pulldowns to further test the effects of *alcA*-Arp1 and *alcA*-Arp11 in dynactin integrity (Figure 4E-4H). We found that loss of Arp1 (*alcA*-Arp1) reduced the amount of p150 pulled down with p50-GFP (Figure 4E, 4F), while it moderately reduced the level of p50-GFP itself (Figure 4E, 4G). Thus, loss of Arp1 affects the assembly of the shoulder complex that includes p150 and p50. No Arp11 was detected in p50-GFP pulldown upon loss of Arp1 (Figure 4E) in three pulldown experiments. Thus, although a Arp11-p50 direct interaction was detected in a yeast two-hybrid assay (Clark and Rose, 2006), the Arp11-p50 connection is mainly mediated by the Arp1 mini-filament in vivo, consistent with a vertebrate dynactin structure (Urnavicius et al., 2015). Interestingly, loss of Arp11 (*alcA*-Arp11) significantly reduced the amount of Arp1 pulled down with p50-GFP, while it had no significant effect on the amount of p150 pulled down with p50-GFP (Figure 4E, 4H). Thus, while Arp11 affects the Arp1 mini-filament significantly, a defective or shortened Arp1 mini-filament may be sufficient for supporting shoulder assembly.

## Discussion

The dynactin complex is critical for cytoplasmic dynein-mediated intracellular transport, but its assembly process remains unclear. Here we have identified VezA as a cellular factor involved in dynactin assembly in *A. nidulans*. We also showed that different parts of dynactin must be assembled in a highly coordinated manner because they affect each other significantly (Figure 4I), and thus, VezA may help coordinate the process to ensure a proper assembly of the complex.

VezA does not affect the assembly of the pointed-end sub-complex, but it affects the assembly of the pointed-end sub-complex onto dynactin, as suggested by results that Arp11-GFP pulls down normal amounts of p25 and p62 but much lower amounts of Arp1, p50, p150 and capping protein in the Δ*vezA* extract (Figure 3F). It is also possible that VezA affects other parts of dynactin. For example, VezA may promote the assembly of the Arp1 mini-filament, and a shortened Arp1 mini-filament in the Δ*vezA* mutant may be defective in its interaction with the dynactin shoulder, thereby causing a decrease in the amounts p150 and p50 pulled down with Arp11-GFP. It is also possible that VezA may affect the assembly of the shoulder, which in turn causes a defect in Arp1 mini-filament integrity. However, a decrease in Arp1 mini-filament length alone in the Δ*vezA* mutant does not explain our data that the capping protein pulled-down with Arp11-GFP is also decreased significantly in the Δ*vezA* extract (Figure 3F).

Except for the pointed-end sub-complex, different parts of the dynactin complex all seem to depend on other parts for integrity (Figure 4I). This dependency most likely happens during dynactin assembly. For example, although the p150-p50-p24 shoulder complex can be separated from the rest of the purified dynactin complex (Eckley et al., 1999), studies from genetic model organisms all showed that p150 stability is compromised upon loss of Arp1 (Haghnia et al., 2007; Minke et al., 1999; Zhang et al., 2008). Our current data show that loss of Arp1 decreases the protein level of p50-GFP and lowers the amount of p150 pulled down with p50-GFP (Figure 4E-4G). Thus, the Arp1 mini-filament is required for the correct assembly of the shoulder containing p50, p24 and the C-terminal parts of p150. Combined with previous data that loss of p24 disrupts the p150-p50 interaction in budding yeast (Amaro et al., 2008) and lowers p150 stability in *N. crassa* (Minke et al., 1999), we think that loss of Arp1 may affect the assembly of the two p50-p24 parts of the shoulder that bind the C-terminal parts of p150 (Urnavicius et al., 2015), thereby affecting p150 stability. Our results also suggest that the shoulder is important for the integrity of the Arp1 mini-filament and/or its connection with the pointed-end sub-complex. Specifically, we found that the amount of Arp1 pulled down with p25-GFP is significantly reduced upon loss of p150 or p50 (Figure 4A-4D). The effect of p150 on Arp1 is likely mediated by p50 as the amount of p50 pulled down with p25-GFP is dramatically reduced in the *alcA*-p150 mutant (Figure 4C, 4D). There are four copies of p50 within dynactin (Eckley et al., 1999), which extend out tentacle-like sites that touch the Arp1 mini-filament (Cheong et al., 2014; Urnavicius et al., 2015). Based on a cryo-EM structure of dynactin, it was proposed that the four p50 subunits and their extended regions may control the Arp1 mini-filament assembly (Urnavicius et al., 2015). Our pulldown result obtained using p25-GFP in the *alcA*-p50 extract is consistent with a shortened Arp1 mini-filament although it can also be interpreted by a weakened connection between Arp1 and the pointed-end sub-complex upon loss of p50. Finally, although purified Arp1 can self-assemble to form a filament in vitro (Bingham and Schroer, 1999), the pointed end plays a significant role in Arp1-mini-filament integrity in vivo. In *A. nidulans*, the amount of Arp1 pulled down with p150 was significantly lowered upon loss of Arp11 or p62 (Zhang et al., 2008), and loss of pointed end severely affected dynactin integrity in mammalian cells (Yeh et al., 2012). In this study, we also showed that loss of Arp11 significantly reduced the amount of Arp1 pulled down with p50-GFP, consistent with a role of the pointed-end sub-complex in promoting Arp1 mini-filament assembly in vivo. Interestingly, loss of Arp11, unlike loss of Arp1, does not significantly affect shoulder assembly (indicated by an association between p50 and p150) (Figure 4E, 4F, 4H), suggesting that a partial assembly of the Arp1 mini-filament is sufficient for shoulder assembly. Together, these data all suggest that the assembly of the dynactin complex needs to be highly coordinated. It is possible that VezA is involved in helping the coordination.

Although VezA could possibly affect multiple parts of dynactin, our VezA-dynactin interaction data suggest that the sites of VezA action are most likely the Arp1 mini-filament and its pointed-end. Specifically, both Arp1 and its pointed end, but not p50 or p150 (the shoulder), are important for the VezA-dynactin interaction. Upon loss of p50, even though the Arp1 mini-filament could possibly be shorter as p25-GFP pulls down lower amount of Arp1 (Figure 4A, 4B), the VezA-Arp1 interaction is not reduced (Figure 2B, 2D). Upon loss of Arp1, the pointed-end proteins were still pulled down with ΔTM-VezA-GFP, although the amounts were reduced compared to when Arp1 is present (Figure 2E, 2F; Table S1). Thus, the pointed-end sub-complex itself is able to interact with VezA. Since VezA is not required for assembly of the pointed-end sub-complex, it is most likely that VezA promotes or stabilizes the connection between Arp1 and the pointed-end sub-complex. The pointed end is critical for cargo binding (Chaaban and Carter, 2022; Drerup et al., 2017; Gama et al., 2017; Lau et al., 2021; Qiu et al., 2018; Urnavicius et al., 2018; Urnavicius et al., 2015; Yeh et al., 2013; Yeh et al., 2012; Zhang et al., 2014; Zhang et al., 2011). Could VezA be required for stabilizing the pointed end during cargo binding? We do not have data to exclude this possibility, but since the microtubule plus-end-accumulation of the pointed-end proteins is dramatically decreased in the Δ*hookA* mutant, VezA is most likely needed before cargo adapter binding.

One limitation of this study is that we have not obtained purified dynactin or the Arp1 mini-filament with its pointed-end sub-complex from *A. nidulans* to test if they bind VezA directly. This has also prevented any structural studies on the interaction of VezA with dynactin or with the Arp1 mini-filament containing its pointed end. However, our AlphaFold2-based prediction suggests that a direct VezA interaction with the Arp1 mini-filament containing its pointed end is possible (Figure S1). The interaction may be transient since VezA may leave dynactin after its role as an assembly factor is accomplished. Could the function of VezA in dynactin assembly be evolutionarily conserved? Recent results from Drosophila and zebrafish suggest that vezatin-like proteins play a conserved role in dynein function during axonal transport of cargoes (Spinner et al., 2020). In addition, two independent lines of evidence suggest that mammalian vezatin (VezA homolog) also interacts with Arp1: First, vezatin (gene name: VEZT) associates with Arp1 in a human interactome study (Hein et al., 2015) (see Table 4 of our previous publication (Qiu et al., 2020)). Second, vezatin is in close proximity to Arp1 as revealed by a human BioID study (https://humancellmap.org/explore/reports/prey?id=VEZT) (Go et al., 2021). Thus, it is possible that vezatin in higher eukaryotic cells plays a similar role in dynactin assembly, but future work will be needed to test this possibility. While VezA is the first identified cellular factor involved in dynactin assembly, it may not be the only one given that the assembly of the dynactin complex needs to be highly coordinated. Dynactin and dynein are both multiple-protein complexes (Rao and Gennerich, 2024; Schroer, 2004), and how multiple-protein complexes are assembled in a coordinated manner is a general question in cell biology. Vezatin and its homologs including VezA contain predicted transmembrane domains as well as disordered regions, and they may associate with vesicles (Spinner et al., 2020; Yao et al., 2015) or ER membrane (https://humancellmap.org/explore/reports/prey?id=VEZT) (Go et al., 2021). Given that vesicle/organelle-bound mRNAs/ribosomes may support local protein translation, function and/or assembly of multi-protein complexes (Baumann et al., 2014; De Pace et al., 2024; Higuchi et al., 2014; Morita et al., 2024; Zander et al., 2016), a possibility worth considering is that dynactin components are translated near VezA/vezatin to be properly assembled into a functional complex.

## Materials and Methods

### *A. nidulans* strains and media

*A. nidulans* strains used in this study are listed in Supplemental Table 3 (Table S3). Solid rich medium was made of either YAG (0.5% yeast extract and 2% glucose with 2% agar) or YAG+UU (YAG plus 0.12% uridine and 0.11% uracil). Genetic crosses were done by standard methods. Solid minimal medium containing 1% glucose was used for selecting progeny from a cross and for selecting diploids. For live-cell imaging, cells were cultured in liquid minimal medium containing 1% glycerol for overnight at 32°C. All the biochemical analyses and genomic DNA preparation were done using cells grown at 32°C for overnight in liquid YG rich medium (0.5% yeast extract and 2% glucose). For experiments using the *alcA-*based conditional-null mutants, we harvested spores from the solid minimal medium containing 1% glycerol and cultured them in rich medium containing glucose, which is repressive for the *alcA* promoter.

### Live-cell imaging and analyses

Microscopic images used in Figure 3 and Figure S1 were generated using a Nikon Ti2-E inverted microscope with Ti2-LAPP motorized TIRF module and a CFI apochromat TIRF 100 x 1.49 N.A. objective lens (oil). The microscope was controlled by NIS-Elements software using 488 nm and 561 nm lines of LUN-F laser engine and ORCA-Fusion BT cameras (Hamamatsu). Images used in Figure 1, Figure S2 and Figure S3 were captured using an Olympus IX73 inverted fluorescence microscope linked to a PCO/Cooke Corporation Sensicam QE cooled CCD camera. This system also includes a UPlanSApo 100x objective lens (oil) with a 1.40 numerical aperture, a filter-wheel system with GFP/mCherry-ET Sputtered series with high transmission (Biovision Technologies), and the IPLab software used for image acquisition and analysis. Image labeling was done using Adobe Photoshop. For all images, cells were grown at 32°C for ∼18 hours in the LabTek Chambered #1.0 borosilicate coverglass system (Nalge Nunc International, Rochester, NY). Images were taken at room temperature. All the images were taken with a 0.1-s exposure time (binning: 2x2). Quantitation of signal intensity was done as described previously (Zhang et al., 2014). Specifically, a region of interest (ROI) was selected and maximal intensity within the ROI was measured. Then the ROI box was dragged outside of the cell to take the background value, which was then subtracted from the intensity value.

Hyphae were chosen randomly from images acquired under the same experimental conditions. For measuring the signal intensity of microtubule plus-end comets formed by GFP-labeled dynein or dynactin components, only the comet closest to hyphal tip was measured. For measuring GFP-dynein signal intensity at septa, usually only the septum most proximal to the hyphal tip was measured.

### Construction of a strain containing the *gpdA*-ΔTM-*vezA*-GFP allele at the *vezA* locus

Strains were constructed by using standard procedures used in *A. nidulans* (Nayak et al., 2006; Szewczyk et al., 2006; Yang et al., 2004). For constructing the *gpdA*-ΔTM-*vezA*-GFP fusion, we first performed PCRs using genomic DNA from XY167 (Yao et al., 2015) and fours oligoes VTMNN (5’-TTCAGGGACACGCATTTGTG-3’), VTMNC (5’-TATTTGCGTGAGACCAGAGC-3’), VTMATGF (5’-ATGGAATCCCTGGTTTACGAGAA-3’) and VTMCC (5’-CGGTCTTCTTCGTTGAGGAC -3’). These PCRs generated two ∼1 kb genomic fragments upstream and downstream of *vezA* translation start site. Using RQ247 (Qiu et al., 2019) genomic DNA and two oligoes, GPDFVTM (5’-GCTCTGGTCTCACGCAAATAGACTCGAGTACCATTTAATTCTATTTGTG-3’) and GPDRVTM (5’-CTCGTAAACCAGGGATTCCATTGTGATGTCTGCTCAAGCGG-3’), we amplified the ∼1.23-kb fragment containing the *gpdA* promoter. We then used VTMNN1 (5’-GCCCTTGCTAGTCGAAGCA-3’) and VTMNC as primers to perform a fusion PCR of the three fragments and generated a ∼3.3 kb fragment containing the *gpdA* promoter upstream of the *vezA* gene. By using VGORFFi (5’-CATACAGGAGGTGGAACTGGTA-3’) and VGUTRRi (5’-AACGACATCAGGAGAGTCGTC-3’) as primers, a ∼4.7 kb fragment containing the C-terminus of VezA linked to GFP and the selectable marker *AfpyrG* (*Aspergillus fumigatus pyrG*) was amplified by PCR from the genomic DNA of XY163 (Yao et al., 2015). The ∼3.3 kb and ∼4.7 kb fragments were co-transformed into the XY42 strain (Qiu et al., 2018) containing Δ*nkuA* (Nayak et al., 2006) and mCherry-RabA (Abenza et al., 2009; Zhang et al., 2010). The primers VGORFF, VGORFFi, VGUTRR, and VGUTRRi have all been described previously (Yao et al., 2015).

For transformation, spores from XY42 were cultured in a flask containing 50 mL YG+UU liquid medium, which was shaken overnight at 80 rpm at room temperature and then at 180 rpm at 32°C for about 1.5 hours. The medium was poured off, and hyphae were then treated with about 20 mL solution containing cell-wall-lysing enzymes. This solution contains 10 mL of solution 1 (52.8 g of ammonium sulfate and 9.6 g of citric acid in 500 mL water, pH adjusted to 6.0 with 5 M KOH), 10 mL of solution 2 (5 g of yeast extract and 10 g of sucrose in 500 mL water), 0.25 mL of 1 M MgSO4, 200 mg of fraction V bovine serum albumin, 0.8 g of Extralyse (Laffort CS61611), and 0.05 mL of β-glucuronidase (Sigma, G8885). This mixture was made and filter-sterilized within 1 hour before being used to treat the hyphae. After about 3 hours of treatment with this solution at 32°C with shaking at 180 rpm, protoplasts were generated. The protoplasts were collected by centrifugation at 1700 rpm for 1 minute using a swing-bucket rotor (Eppendorf S-4-72), washed with 15 mL of ice-cold solution 3 (26.4 g of ammonium sulfate, 5 g of sucrose and 4.8 g of citric acid in 500 mL water, pH adjusted to 6.0 with 5 M KOH), and finally suspended in 0.5 mL of ice-cold solution 5 (4.48 g of KCl, 0.75 g of CaCl_2_ and 0.195 g of MES in 100 mL water, pH adjusted to 6.0 with 5 M KOH). In a 15-mL tube, 100 μL protoplast was mixed with 20 μL DNA (1 – 2 μg total) and 50 μL ice-cold solution 4 (25 g of PEG 6000 or 8000 (Sigma, P2139), 1.47 g of CaCl_2_.2H_2_O, 4.48 g of KCl, and 1.0 mL of 1 M Tris-HCl pH 7.5 in 100 mL water). This mixture was kept on ice for 20 min, followed by addition of 1 mL solution 4 with gentle mixing. The tube was kept at room temperature for 20 minutes. 10 mL of 50°C pre-melted solid medium (YAG + 0.6 M KCl) was added into the tube and the mixture was poured into a petri dish with a thin layer of the same solid medium (YAG + 0.6 M KCl). After the plates were incubated at 37°C for 2-3 days, colonies of transformants appeared. Autoclaved toothpicks were used to touch the top of the individual colony and transfer the asexual spores onto a YAG plate, which was incubated at 37°C for 2 days. The transformants were then screened by microscopically observing the GFP signals, and the presence of ΔTM-VezA-GFP was confirmed by western blotting analysis with a mouse monoclonal anti-GFP antibody from Clontech. In addition, we also performed a diagnostic PCR to verify the homologous integration of the ∼3.3-kb fragment using VTMNN and GPDRVTM. We also verified the homologous integration of the ∼4.7-kb fragment using AfpyrG5 (5’-AGCAAAGTGGACTGATAGC-3’) and VGUTRR (5’-AGTGCTCCTGGTCAATGTCCA-3’).

### Construction of the p150 conditional-null mutant, *alcA*-p150

In the *alcA*-p150 strain, the *alcA* promoter is inserted in front of the *nudM* gene encoding dynactin p150 to conditionally shut off its expression. This was done by first making a ∼2.4 kb fragment containing the *AfpyrG* selectable marker linked upstream to the *alcA*-promoter (AfpyrG-alcA). Specifically, we used ALCAR (5’-TTTGAGGCGAGGTGATAGGA-3’) and ALCAF (5’-AGACCGAGTGAACGTATACC-3’) as primers to amplify a ∼0.5 kb alcA fragment from GR5 genomic DNA. We then used APYRGR (5’-GGTATACGTTCACTCGGTCTCTGTCTGAGAGGAGGCACTGA-3’) and APYRGF (5’-TGCTCTTCACCCTCTTCGCG-3’) to amplify a ∼1.9 kb AfpyrG fragment from the plasmid pAO81. We then fused these two fragments by using APYRGF and ALCAR as primers to obtain the ∼2.4 kb AfpyrG-alcA fragment. To insert this ∼2.4 kb AfpyrG-alcA fragment upstream of the p150 gene, we used the following six oligoes: p150NN (5’-ATCTGTAAGGGTCGCACGG-3’), p150NN1 (5’-TTGTCGCCAGGAGAGCCTG-3’), p150NC (5’-GGTATACGTTCACTCGGTCTTGTTGTTATTGACCGCGCC-3’), p150CN (5’-TCCTATCACCTCGCCTCAAATTCCAAACACAACGGCCGT-3’), p150CC (5’-CTCAATTCGTCAAGCTCCCTG -3’) and p150CC1 (5’-ATTGCCATGTTCTGAGACTGTCG -3’). Specifically, p150NN, p150NC, p150CN and p150CC were used to amplify two ∼1 kb fragments upstream and downstream of the p150 translational start site from GR5 genomic DNA. These fragments were fused to the ∼2.4 kb AfpyrG-alcA fragment by a fusion PCR using p150NN1 and p150CC1 as primers. This 4.4 kb fusion fragment was transformed into the XY42 strain to get the *alcA*-p150 strain. Transformants that exhibited a compact-colony phenotype on the glucose-containing YAG medium were selected for PCR verification of the correct genotype. Specifically, genomic DNA was extracted and PCR reactions were performed to confirm the homologous integration event using alcA5 (5’-AGCACTTTCTGGTACTGTCC-3’) and p150CC as primers.

### Construction of the p50 conditional-null mutant, *alcA*-p50

In the *alcA*-p50 strain, the *alcA* promoter is inserted in front of the p50 gene to conditionally shut off its expression. This was done by first making a ∼2.4 kb fragment containing the *AfpyrG* selectable marker linked upstream to the *alcA*-promoter (AfpyrG-alcA), as described above. To insert this ∼2.4 kb AfpyrG-alcA fragment upstream of the p50 gene, we used the following six oligoes: p50NN (5’-AGGGAGGTTTGAACCATGG -3’), p50NC (5’-CGCGAAGAGGGTGAAGAGCAAGCTAGAATATTGAAGGATCTTAGTTGTC -3’), p50CN (5’-TCCTATCACCTCGCCTCAAAATGGCTTTCAACAAAAAATATGCTGGTC -3’), p50CC (5’-AAGTGCTTCAGCGTCTGCTG -3’), p50NN1 (5’-TCGGAGATGGTTCGATCCTG -3’) and p50CC1 (5’-TTCCTGACGCGGGTAGAAAG -3’). Specifically, p50NN, p50NC, p50CN and p50CC were used to amplify two ∼1 kb fragments upstream and downstream of the p50 translational start site from GR5 genomic DNA. These fragments were fused to the ∼2.4 kb AfpyrG-alcA fragment by a fusion PCR using p50NN1 and p50CC1 as primers. This 4.4 kb fusion fragment was transformed into the XY42 strain to get the *alcA*-p50 strain. Transformants that exhibited a compact-colony phenotype on the glucose-containing YAG medium were selected for PCR verification of the correct genotype. Specifically, genomic DNA was extracted and PCR reactions were performed to confirm the homologous integration event using two pairs of primers: (1) alcA5 and p50CC; (2) AfpyrG3 (5’-GTTGCCAGGTGAGGGTATTT-3’) and p50NN.

### Construction of the Arp1 conditional-null mutant, *alcA*-Arp1

In the *alcA*-Arp1 strain, the *alcA* promoter is inserted in front of the *nudK* (Arp1) gene to conditionally shut off its expression. This was done by first making a ∼2.4 kb fragment containing the *AfpyrG* selectable marker linked upstream to the *alcA*-promoter (AfpyrG-alcA), as described above. To insert this ∼2.4 kb AfpyrG-alcA fragment upstream of the Arp1 gene, we used the following six oligoes: Arp1NN (5’-TGGCAAGGACGGACAGCAG -3’), Arp1NC (5’-CGCGAAGAGGGTGAAGAGCATGCAGGGAATTGGTTGCGAG -3’), Arp1CN (5’-TCCTATCACCTCGCCTCAAAATGACCGAGGCTACTCTTCAC -3’), Arp1CC3 (5’-AATGTTGAGGTAAAGCGACTTGCG -3’), Arp1NN1 (5’-TTTGACGACTTTGTCGCAAAC -3’) and Arp1CC2 (5’- CAAGTCGAGATCCGTAGGAT -3’). Specifically, Arp1NN, Arp1NC, Arp1CN and Arp1CC3 were used to amplify two ∼1 kb fragments upstream and downstream of the Arp1 translational start site from GR5 genomic DNA. These fragments were fused to the ∼2.4 kb AfpyrG-alcA fragment by a fusion PCR using Arp1NN1 and Arp1CC2 as primers. This 4.35 kb fusion fragment was transformed into XY42 and RQ54 strains to get *alcA*-Arp1 strains. Transformants that exhibited a compact-colony phenotype on the glucose-containing YAG medium were selected for PCR verification of the correct genotype. Specifically, genomic DNA was extracted and PCR reactions were performed to confirm the homologous integration event using two pairs of primers: (1) alcA5 and Arp1CC3; (2) AfpyrG3 and Arp1NN.

### Construction of the Arp11 conditional-null mutant, *alcA*-Arp11

In the *alcA*-Arp11 strain, the *alcA* promoter is inserted in front of the Arp11 gene to conditionally shut off its expression. This was done by first making a ∼2.4 kb fragment containing the *AfpyrG* selectable marker linked upstream to the *alcA*-promoter (AfpyrG-alcA), as described above. To insert this ∼2.4 kb AfpyrG-alcA fragment upstream of the Arp11 gene, we used the following six oligoes: Arp11NN (5’- CAGCTTCTTTCGAGAAGTTCATG -3’), Arp11NC (5’- CGCGAAGAGGGTGAAGAGCATCTCGTCAATAAATTTCTGTGGTTGG -3’); Arp11CN (5’- TCCTATCACCTCGCCTCAAAATGTCCTCAATGTCGATCCGC -3’), Arp11CC (5’- GTATACCAGTAGAGAGATCGGC -3’), Arp11NN1 (5’- CAGTAAATCCCGATCCAAACATCAG - 3’) and Arp11CC1 (5’- TTCTTGGTCGTCCAGGTCG -3’). Specifically, Arp11NN, Arp11NC, Arp11CN and Arp11CC were used to amplify two ∼1 kb fragments upstream and downstream of the Arp11 translational start site from GR5 genomic DNA. These fragments were fused to the ∼2.4 kb AfpyrG-alcA fragment by a fusion PCR using Arp11NN1 and Arp11CC1 as primers. This 4.35 kb fusion fragment was transformed into the XY42 strain to get the *alcA*-Arp11 strain. Transformants that exhibited a compact-colony phenotype on the glucose-containing YAG medium were selected for PCR verification of the correct genotype. Specifically, genomic DNA was extracted and PCR reactions were performed to confirm the homologous integration event using two pairs of primers: (1) AfpyrG3 and Arp11NN; (2) alcA5 and Arp11CC. We note that the conditional-null mutants including *alcA*-p150, *alcA*-p50, *alcA*-Arp1 and *alcA*- Arp11 constructed in this study all contain the *alcA*-driven gene integrated as a linear-fragment replacing the endogenous gene, which is more stable than the circular plasmid-based integration in previously constructed *alcA* mutants (Zhang et al., 2008).

### Construction of the *vezA*^Δ1-20^-GFP mutant allele at the *vezA* locus

In the *vezA*^Δ1-20^-GFP strain, the sequence encoding the first 20 amino acids of VezA is deleted, and GFP is linked to the C-terminus of VezA. This was done by first making a ∼2.0 kb fragment of vezA^Δ1-20^. Specifically, by using GR5 genomic DNA as template, a 1.1 kb and 1.0 kb PCR products were obtained using two pairs of primers: (1) VTMNN (5’- TTCAGGGACACGCATTTGTG-3’) and VNDNC (5’-GCCAGTCTGAACTATGTTCACCCATAATTTATGCTCATTCCGTAAGCG -3’), (2) VNDCN (5’- GGTGAACATAGTTCAGACTGGC -3’) and VTMCC (5’- CGGTCTTCTTCGTTGAGGAC-3’). These two fragments were used to make a fusion PCR product of ∼2.0 kb by using primers VTMNN1 (5’-GCCCTTGCTAGTCGAAGCA-3’) and VTMCC. By using VGORFFi (5’-CATACAGGAGGTGGAACTGGTA-3’) and VGUTRRi (5’- AACGACATCAGGAGAGTCGTC-3’) as primers, a ∼4.7 kb fragment containing the C-terminus of VezA linked to GFP and the selectable marker *AfpyrG* was amplified by PCR from the genomic DNA of XY163 (containing *vezA*-GFP) (Yao et al., 2015). The ∼2.0 kb and ∼4.7 kb fragments were co-transformed into the RQ54 strain. Transformants were examined microscopically, and those with an abnormal early-endosome accumulation at the hyphal tip were selected, and the correct genotype was confirmed by PCR with the primers AfpyrG5 (5’- AGCAAAGTGGACTGATAGC-3’) and VGUTRR (5’-AGTGCTCCTGGTCAATGTCCA-3’) followed by sequencing of the PCR product using primers VNDSq5 (5’ CTCACGTCACGACTTGTCGA-3’), VTMATGF (5’-ATGGAATCCCTGGTTTACGAGAA-3’) and GFP5R (5’-CCAGTGAAAAGTTCTTCTCCTTTAC-3’).

### Construction of the *vezA*^Δ563-615^-GFP mutant allele at the *vezA* locus

In the *vezA*^Δ563-615^-GFP strain, the sequence encoding the last 53 amino acids of VezA is deleted, and GFP is linked to the C-terminus of VezA. This was done by first making a ∼3.8 kb GFP-*AfpyrG*-containing fragment from XY163 genomic DNA using GAGAF (5’- GGAGCTGGTGCAGGCGCTG-3’) and VGUTRR (5’-AGTGCTCCTGGTCAATGTCCA-3’) as primers. We then used TMTNC3 (5’- GGCACCGGCTCCAGCGCCTGCACCAGCTCCAGAAGCTCGCTTGTTATTTCGC-3’) and VGORFF (5’-TCGATGCTGCTGTGCTGTTGA-3’) to obtain a 0.9 kb fragment. The 3.8 kb and the 0.9 kb fragments were fused to obtain a 4.7 kb fragment in a fusion PCR reaction using primers VGORFFi (5’-CATACAGGAGGTGGAACTGGTA-3’) and VGUTRRi (5’- AACGACATCAGGAGAGTCGTC-3’). The 4.7 kb fragment was transformed into the strain XY42. Transformants were examined microscopically, and those with an abnormal early-endosome accumulation at the hyphal tip were selected, and the correct genotype was confirmed by PCR with the primers AfpyrG5 and VGUTRR followed by sequencing of the PCR product using primers VTMATGF and GFP5R.

### Construction of the p50-GFP and p50-S alleles at the p50 gene locus

Eight primers were used to make the p50-GFP strain: P50GNN (5’- AAGATGAGATGGCGGCGTC-3’), P50GNN1 (5’-CGAGGCGAAGGACACATCA-3’), P50GNC (5’-GCTCCAGCGCCTGCACCAGCTCCCTTCCCACTCTCCAACTTCTCC-3’), P50GCC (5’- TCCAACCACACAAGGAATG -3’), P50GCC1 (5’-CATCTTTAACAGCTGCCGCC-3’), P50GCN (5’-CATCAGTGCCTCCTCTCAGACAGAGTACTTAATATAGTGTAAGGTGAGATG-3’), new pyrG3 (5’-CTGTCTGAGAGGAGGCACTGATGCG-3’) and GAGAF (5’- GGAGCTGGTGCAGGCGCTG-3’). Specifically, P50GNN and P50GNC were used to obtain a ∼1 kb p50 open-reading frame fragment using GR5 wild-type strain genomic DNA as template. P50GCC and P50GCN were used to obtain a ∼1 kb fragment covering the 3’ untranslated region of the gene using GR5 genomic DNA as template. GAGAF and new pyrG3 were used to obtain a 3.6 kb GFP-AfpyrG fragment using the pFNO3 plasmid as template. Primers P50GNN1 and P50GCC1 were used to fuse the three fragments together by fusion PCR to obtain a 4.6 kb fragment, which was transformed into the RQ54 strain. Transformants were screened for GFP signals under microscope, and homologous integration was confirmed by PCR using AfpyrG5 and P50GCC as primers.

The same eight primers were also used for making the p50-S strain. Both franking fragments were the same as above, but GAGAF and new pyrG3 were used to obtain a 3.0 kb S-AfpyrG fragment using the pAO81 plasmid as template. Primers P50GNN1 and P50GCC1 were used to fuse the three fragments together by fusion PCR to obtain a 4.0 kb fragment, which was transformed into the XY42 strain. The selected transformant was confirmed by PCR using AfpyrG5 and P50GCC as primers and by western blot analysis.

### Construction of a strain containing the p62-GFP allele at the p62 gene locus

For constructing the p62-GFP fusion, we used the following six primers to amplify wild type genomic DNA and the GFP-AfpyrG fusion from the plasmid pFNO3: P62GNC (5’- GGCTCCAGCGCCTGCACCAGCTCCTGAAGAACTGGAACTGCCAGC-3’), P62GNN (5’- CGAGTCTGAAGTTGCCATCATC-3’), P62GCN (5’- ATCAGTGCCTCCTCTCAGACAGTTCTCCTTCACCCTTATCTACTATATTC -3’), P62GCC (5’- GCATTGTTGTTAGGCAGTGGC -3’), GAGAF (5’- GGAGCTGGTGCAGGCGCTG -3’) and pyrG3 (5’- CTGTCTGAGAGGAGGCACTGAT-3’). A fusion PCR was performed using P62GNN and P62GCC as primers to generate the 4.7 kb P62-GFP-AfpyrG fragment that we used to transform into XY42. Transformants were screened for GFP signals under microscope, and homologous integration was confirmed by PCR using primers AfpyrG5 (5’- AGCAAAGTGGACTGATAGC-3’) and P62GCC2 (5’-TGGCAGCGAATGGAGGCATT-3’).

### Construction of a strain containing the p25-S allele at the p25 gene locus

For constructing the p25-S fusion, we used the following six primers to amplify p25 open reading frame from the XY41 strain and the S-AfpyrG fusion from the plasmid pAO81: ORFF (5’-TATGAGCTTAGCCTGCCCCAC-3’), ORFR (5’-TCGGATACTTCGATATCTCTCCCG-3’), FUSF (5’-TCGGGAGAGATATCGAAGTATCCGAGGAGCTGGTGCAGGCGCTGGAG-3’), FUSR (5’- CCGAGGCCGACTCCAAGTACAGTACCTGTCTGAGAGGAGGCACTGATG-3’), UTRF (5’- GTACTGTACTTGGAGTCGGCCTCG-3’) and Q6 (5’- CGAATCTTCAACTCCTGGGTGCG-3’). A fusion PCR was performed using ORFF2 (5’- AGTAAGTGTACTGTGGCTTCACCG-3’) and UTRR2 (5’-GTCATTGACTATCCTGCAGGTGAG- 3’) as primers to generate the 3.9 kb p25-S-AfpyrG fragment that we used to transform into RQ54. Transformants were screened for homologous integration by PCR using Q2 (5’- TGGTAATAGACGGCAGTGGG-3’) and STAG3 (5’-GCTGGCGTTCGAATTTAGC-3’) as primers.

### Biochemical pull-down assays, western analysis and antibodies

The μMACS GFP-tagged protein isolation kit (Miltenyi Biotec) was used to pull down proteins associated with the GFP-tagged protein. This was done as described previously (Zhang et al., 2014). Specifically, about 0.6 g hyphal mass was harvested from overnight culture for each sample, and cell extracts were prepared using a lysis buffer containing 50 mM Tris-HCl (pH 8.0), 0.01% Triton X100 and 10 μg/mL of a protease inhibitor cocktail (Sigma-Aldrich). Cell extracts were centrifuged at 20,000 *g* for 60 minutes at 4°C, and supernatant was used for the pull-down experiment. To pull down GFP-tagged proteins, 35 μL anti-GFP MicroBeads were added into the cell extracts for each sample and incubated at 4°C for 60 minutes. The MicroBeads/cell extracts mixture was then applied to the μColumn followed by gentle wash three times with the lysis buffer used above for protein extraction (Miltenyi Biotec). Pre-heated (95°C) SDS-PAGE sample buffer was used as elution buffer. Western blot analyses were performed using the alkaline phosphatase system and blots were developed using the AP color development reagents from Bio-Rad. Quantitation of the protein band intensity was done using the Image Studio Lite software (version 5.2). Specifically, an area containing the whole band was selected as a region of interest, and the intensity sum within the region of interest was measured. Then, the region of interest box was dragged to the equivalent region of the negative control lane or a blank region without any band on the same blot to take the background value, which was then subtracted from the intensity sum. The rabbit polyclonal antibody against GFP was from Takara Bio Inc (Catalog number: 632592). The rabbit monoclonal antibody against the S-tag was from Cell Signaling Technology (Catalog number: 12774S). Polyclonal antibody against dynactin p150 was generated in a previous study by injecting proteins produced in bacteria into rabbits followed by affinity purification of the antibody (Zhang et al., 2008). Polyclonal antibody against Arp1 was generated using the service of Pacific Immunology (www.pacificimmunology.com). An immunograde peptide of Cys-SADEWHEDPEIIHRKFA (Arp1 amino acids 364-380) was synthesized and conjugated to the KLH carrier protein, and this conjugated form was used as an antigen for rabbit injection. The antibody was affinity-purified using the same peptide. Polyclonal antibody against Arp11 was generated using the service of Boster Antibody and Elisa experts (www.bosterbio.com). A recombinant protein of 217 amino acids (RSALVVDIGWAETVVSGIYEYREVTTKRSTRAMRSLIQETGRMFTRLLGGDSQPDTISVEFEFCEEVVSRFAWCQPSRSGYYKETAENSLADILDKTISIPSPSNPGSSDIELPFSKLEELVEKVLLAQGMADSDLDDQEKPISLLVYNTLLSLPPDVRGICMSRIVFVGGGANIAGIRSRILDEVAHLIELYGWSPVRGRLIEQQIQKLQSLKLSQ) was used for rabbit injection and purification of serum.

### Statistical analysis

All statistical analyses were done using GraphPad Prism 10 for macOS (version 10.1.1, 2023). The D’Agostino & Pearson normality test was performed on all data sets except for data sets with small n (n=3). For western blot data, mass spectrometry data and the percentage of hyphal tips with abnormal accumulation of early endosomes, data distribution was assumed to be normal but this was not formally tested because the n number is 3 for these data sets. For these data, student *t*-test (unpaired, two-tailed) was used for analyzing any two data sets and ordinary one-way ANOVA (unpaired) was used for analyzing multiple data sets. Note that adjusted p values were generated from the ordinary one-way ANOVA test with Dunnett’s multiple comparisons test. The data sets presented in Figures 1B, 1D, 1F, Figure 3C and Figure S2C did not pass the D’Agostino & Pearson normality test (alpha=0.05), and thus, they were analyzed using the Mann-Whitney test (unpaired, two tailed), a non-parametric test without assuming Gaussian distribution. The data set presented in Figure S3C passed the D’Agostino & Pearson normality test, and thus, they were analyzed using the student *t*-test (unpaired, two tailed).

### ColabFold protein structure prediction

Protein structure prediction was done using ColabFold version 1.5.2 (alphaFold2_multimer_V3) in Google Cloud Platform with NVIDIA A100 80GB GPU (Mirdita et al., 2022). For each protein-complex prediction, we set parameters of num_models to 5, num_recycles to 25, recycle_early_stop_tolerance to 0.5, and use_templates to false. The final models were analyzed with UCSF ChimeraX.

## Acknowledgements

We thank Andrew Carter for very helpful discussions, especially for generously sharing unpublished AlphaFold2-based structural predictions on VezA dimerization and a possible vezatin-Arp1 interaction via the C-terminus of vezatin, which stimulated some experiments described in this manuscript. We thank Berl Oakley, Miguel Peñalva, Samara Reck-Peterson and Martin Egan for Aspergillus strains, the Fungal Genetic Stock Center for the pFNO3 and pAO81 plasmids, and Stephen Osmani for depositing the plasmids. We also thank the Student Bioinformatics Initiative (led by Andrew Frank) at the Uniformed Services University of the Health Sciences for providing high-performance computing resources for protein structure prediction. This work was funded by the National Institutes of Health R35GM140792 (to X.X.) and a Department of Defense #OSD(HA).2022ICD.WBH-1 Joint DOTML-PF Warfighter Brain Health grant to USUHS Department of Biochemistry and Molecular Biology. Disclaimer: The opinions and assertions expressed herein are those of the author(s) and do not necessarily reflect the official policy or position of the Uniformed Services University or the Department of Defense.

---

**Figure S1.**
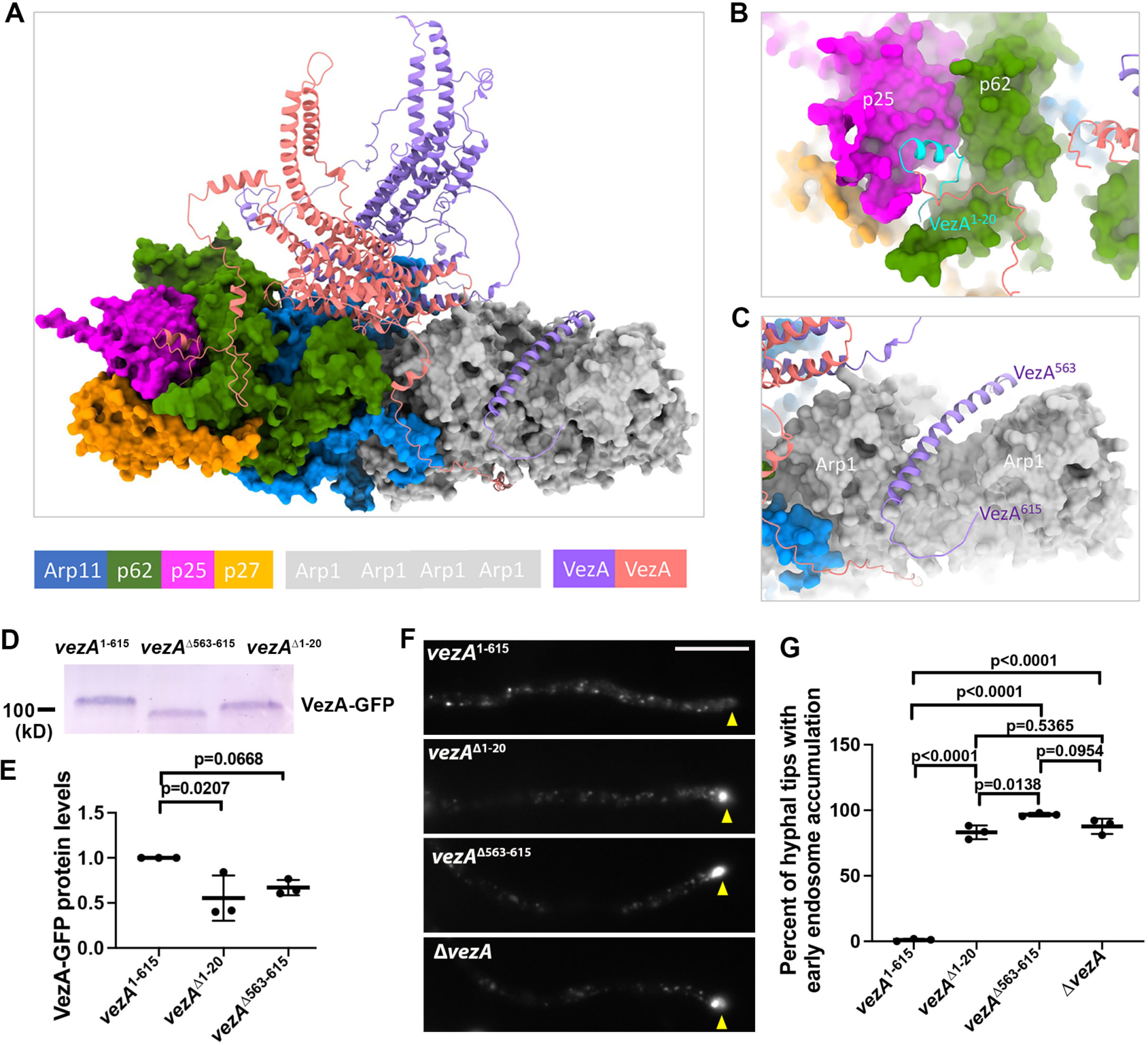
An AlphaFold2-based analysis of VezA-dynactin interaction. (A) An AlphaFold2 prediction model of the VezA dimer binding to Arp1 as well as the pointed end. In this prediction, we included four copies of Arp1 but no convention actin, although conventional actin is located next to Arp11 in the structure of vertebrate dynactin (Urnavicius et al., 2015). This is because our pulldown data were not able to provide evidence that conventional actin is a component of dynactin in *A. nidulans* (Zhang et al., 2018). (B) In this model, the N-terminus of VezA (VezA^1-20^) is close to p25 and p62 of the pointed end. (C) In the same model, the C-terminus of VezA (VezA^563-615^) is close to the Arp1 mini-filament. (D) A western blot showing the protein levels of VezA-GFP in the *vezA*^1-615^-GFP (full-length), *vezA*^Δ1-20^-GFP and *vezA*^Δ563-615^-GFP strains. (E) A quantitative analysis on the effects of the *vezA*^Δ1-20^ and *vezA*^Δ563-615^ mutations on the level of VezA-GFP. The values were generated from western blot analyses of three independent pull-down experiments (*n* = 3 for all). Scatterplots with mean and SD values as well as p values were generated by Prism 10 (ordinary one-way ANOVA test with Dunnett’s multiple comparisons test). (F) Microscopic images showing the distributions of mCherry-RabA-labeled early endosomes in the *vezA*^Δ1-20^ and *vezA*^Δ563-615^ mutants. Hyphal tip is indicated by a yellow arrowhead. Bar, 10 μm. (G) A quantitative analysis on the percentage of hyphal tips with the abnormal accumulation of early endosomes. Three experiments were performed, and in each experiment, 50 or more hyphal tips were examined for each strain. Scatterplots with mean and SD values as well as p values were generated by Prism 10 (ordinary one-way ANOVA test with Tukey’s multiple comparisons test).

**Figure S2.**
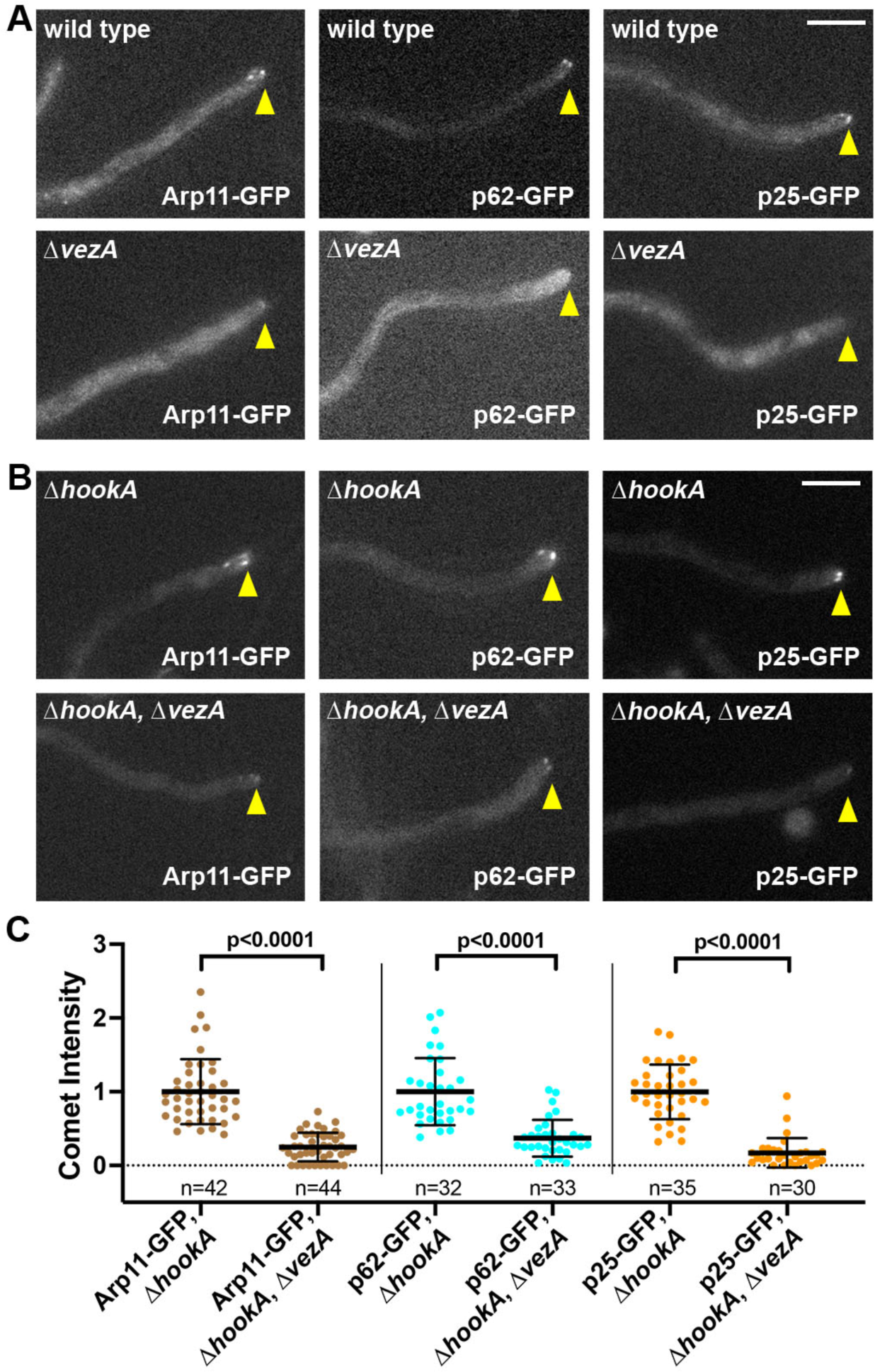
VezA affects the microtubule plus-end localization of several pointed-end proteins. (A) Images of Arp11-GFP, p62-GFP and p25-GFP accumulation at the microtubule plus ends as represented by comets near the hyphal tip in the wild-type and Δ*vezA* strains. Hyphal tip is indicated by a yellow arrowhead. Bar, 5 μm. (B) Images of Arp11-GFP, p62-GFP and p25-GFP in the Δ*hookA* single mutant and the Δ*hookA*, Δ*vezA* double mutant strains. Hyphal tip is indicated by a yellow arrowhead. Bar, 5 μm. (C) Quantitative analyses on plus-end comet intensity of Arp11-GFP, p62-GFP and p25-GFP in the Δ*hookA* and Δ*hookA*, Δ*vezA* strains. The average value for the intensity in the Δ*hookA* single mutant is set as 1. Scatter plots with mean and S.D. values were generated by Prism 10, and the p values were generated by Mann-Whitney test (unpaired).

**Figure S3.**
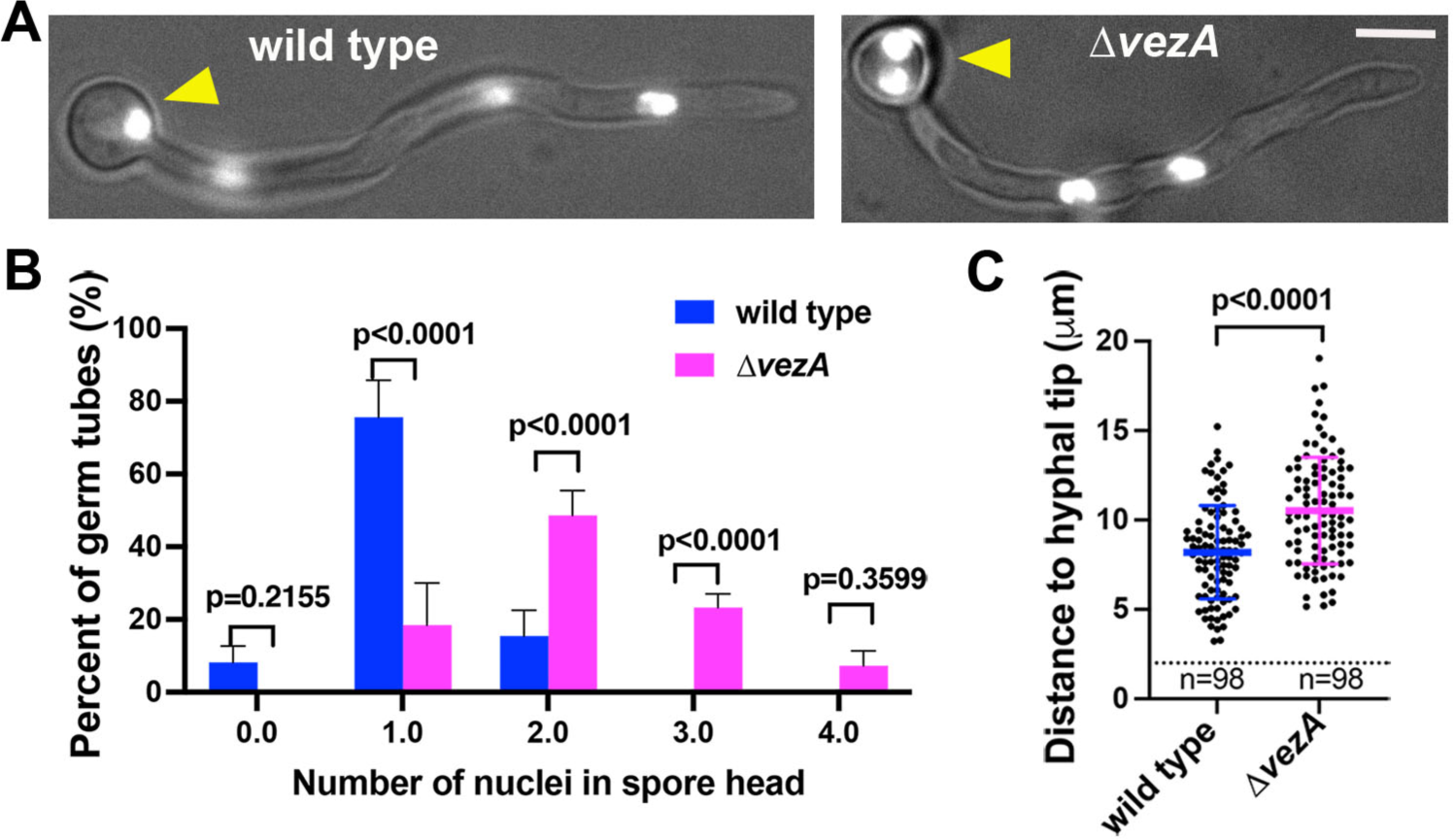
Nuclear distribution is defective in the Δ*vezA* mutant. (A) Histone H1-GFP-labeled nuclei in wild type and the Δ*vezA* mutant. Strains were grown overnight at 32°C in liquid minimal medium containing 1% glycerol. Yellow arrow head indicates the spore swelling. Bar, 5 μm. (B) A quantitative analysis on the percent of germ tubes containing different numbers of nuclei in the spore swelling. Column bar graphs with mean and S.D. values were generated from five experiments. For each experiment, at least 36 germ tubes were counted from each strain, and the total numbers of counted germ tubes are 229 for the wild-type control and 238 for the Δ*vezA* mutant. P values were generated from two-way ANOVA with Bonferroni’s multiple comparisons test. (C) A quantitative analysis on the distance from the hyphal tip to the nucleus closest to it. Scatterplots with mean and SD values as well as p values were generated by Prism 10 (Student’s *t* test, two tailed, unpaired).

**Table S1.** Mass spectrometry data showing numbers of total peptides and unique peptides (in parentheses) of proteins pulled down with ΔTM-VezA-GFP in the wild-type background (Control) and in the *alcA*-Arp11, *alcA*-p50 or *alcA*-Arp1 background where expression of the *alcA*- promoter-controlled gene is repressed by glucose.

| Protein/<br>numbers of total and<br>unique peptides<br>(unique peptides in<br>parentheses) | Control | *alcA*-<br>Arp11 | *alcA*-<br>p50 | *alcA*-<br>Arp1 | *alcA*-<br>Arp11/<br>control<br>(ratio of<br>total<br>peptides) | *alcA*-<br>p50/<br>control<br>(ratio of<br>total<br>peptides) | *alcA*-<br>Arp1/<br>control<br>(ratio of<br>total<br>peptides) |
| --- | --- | --- | --- | --- | --- | --- | --- |
| **Dynein HC** (An0118,<br>4345 aa) | 199<br>(183) | 13<br>(13) | 17<br>(17) | 7<br>(6) | 0.07 | 0.09 | 0.04 |
| **p150** of dynactin<br>(An6323, 1342 aa) | 59<br>(57) | 3<br>(3) | 0<br>(0) | 0<br>(0) | 0.05 | 0 | 0 |
| **p50** of dynactin<br>(An3589, 467 aa) | 24<br>(23) | 2<br>(2) | 1<br>(1) | 0<br>(0) | 0.08 | 0.04 | 0 |
| **Arp1** of dynactin<br>(An1953, 380 aa) | 30<br>(25) | 5<br>(5) | 19<br>(17) | 2<br>(2) | 0.17 | 0.63 | 0.07 |
| **Arp11** of dynactin<br>(An3185, 557 aa) | 34<br>(29) | 1<br>(1) | 31<br>(26) | 12<br>(11) | 0.03 | 0.91 | 0.35 |
| **p62** of dynactin<br>(An4917, 637 aa) | 18<br>(17) | 1<br>(1) | 22<br>(21) | 4<br>(4) | 0.06 | 1.22 | 0.22 |
| **p25** of dynactin<br>(An5022, 202 aa) | 6<br>(6) | 0<br>(0) | 6<br>(6) | 3<br>(3) | 0 | 1.00 | 0.50 |
| **GFP**<br>(238 aa) | 29<br>(17) | 28<br>(12) | 29<br>(16) | 27<br>(12) | 0.97 | 1.00 | 0.93 |
| **Myosin V**<br>(An8862, 1569aa) | 105<br>(94) | 106<br>(75) | 106<br>(84) | 106<br>(85) | 1.01 | 1.01 | 1.01 |
| **NudF/LIS1**<br>(An6197, 444aa) | 18<br>(16) | 3<br>(3) | 2<br>(2) | 0<br>(0) | 0.17 | 0.11 | 0 |

Note that ΔTM-VezA-GFP also pulled down Myosin V but the amount of pulled-down Myosin V is not affected by the loss of Arp11 (*alcA*-Arp11), p50 (*alcA*-p50) or Arp1 (*alcA*-Arp1).

**Table S2.** Mass spectrometry data from three experiments showing numbers of total peptides and unique peptides of proteins pulled down with Arp11-GFP in the wild-type background (*vezA*+) and in the Δ*vezA* background.

| Protein/<br>number of<br>peptides<br>(Experiment 1) | vezA+<br>(unique) | **vezA+**<br>**(total)** | ΔvezA<br>(unique) | **ΔvezA**<br>**(total)** | **ΔvezA/vezA+ (ratio**<br>**of total peptides**<br>**relative to the Arp11**<br>**ratio set as 1)** |
| --- | --- | --- | --- | --- | --- |
| **p150** (An6323,<br>1342 aa) | 40 | **78** | 12 | **19** | **0.24** |
| **p50** (An3589,<br>467 aa) | 21 | **67** | 10 | **14** | **0.21** |
| **Arp1** (An1953,<br>380 aa) | 12 | **79** | 5 | **13** | **0.13** |
| **Arp11** (An3185,<br>557 aa) | 17 | **51** | 19 | **66** | **1** |
| **p62** (An4917,<br>637 aa) | 12 | **24** | 9 | **26** | **0.84** |
| **p25** (An5022,<br>202 aa) | 4 | **12** | 5 | **24** | **1.55** |
| **Cap alpha**<br>(An2126, 273 aa) | 6 | **13** | 4 | **6** | **0.36** |
| **Cap beta**<br>(An0290, 266aa) | 4 | **8** | 0 | **0** | **0** |
| **Dynein HC**<br>(An0118, 4345 aa) | 88 | **150** | 27 | **34** | **0.18** |
| (Experiment 2) |  |  |  |  |  |
| **p150** | 63 | **88** | 12 | **12** | **0.16** |
| **p50** | 20 | **34** | 10 | **11** | **0.37** |
| **Arp1** | 22 | **56** | 11 | **12** | **0.24** |
| **Arp11** | 17 | **32** | 18 | **28** | **1** |
| **p62** | 17 | **22** | 14 | **19** | **0.99** |
| **p25** | 6 | **8** | 6 | **7** | **1** |
| **Cap alpha** | 6 | **9** | 2 | **2** | **0.25** |
| **Cap beta** | 7 | **10** | 1 | **1** | **0.11** |
| **Dynein HC** | 174 | **224** | 85 | **91** | **0.46** |
| (Experiment 3) |  |  |  |  |  |
| **p150** | 103 | **158** | 77 | **144** | **0.31** |
| **p50** | 25 | **51** | 19 | **66** | **0.44** |
| **Arp1** | 20 | **86** | 26 | **60** | **0.24** |
| **Arp11** | 28 | **45** | 32 | **132** | **1** |
| **p62** | 19 | **24** | 27 | **79** | **1.12** |
| **p25** | 7 | **10** | 7 | **29** | **0.99** |
| **Cap alpha** | 9 | **9** | 9 | **19** | **0.72** |
| **Cap beta** | 12 | **15** | 7 | **7** | **0.16** |
| **Dynein HC** | 243 | **479** | 211 | **552** | **0.39** |

**Table S3.** *Aspergillus nidulans* strains used in this study

| Strain | Genotype | Source |
| --- | --- | --- |
| GR5 | *pyrG89; pyroA4; wA3* | G.S. May |
| JZ505 | Δ*hookA*-Afp<sub>pyrG</sub>; GFP-*nudA*<sup>HC</sup>; *argB2::[argB\*-alcAp::mCherry-RabA]*; possibly *pyrG89*; possibly Δ*nkuA::argB*; *pyroA4; wA2* | (Zhang et al., 2014) |
| JZ788 | Arp11-GFP-Afp<sub>pyrG</sub>, *argB2::[argB\*-alcAp::mCherry-RabA]*; Δ*nkuA::argB*; *pyrG89; pantoB100; yA2* | (Qiu et al., 2020) |
| RQ2 | GFP-*nudA*<sup>HC</sup>; *argB2::[argB\*-alcAp::mCherry-RabA]*; Δ*nkuA::argB*; *pyrG89; pyroA4; yA2* | (Qiu et al., 2013) |
| RQ54 | *argB2::[argB\*-alcAp::mCherry-RabA]*; Δ*nkuA::argB*; *pyrG89; pyroA4; wA2* | (Qiu et al., 2013) |
| RQ294 | GFP-*nudA*<sup>R1602E, K1645E</sup>; *argB2::[argB\*-alcAp::mCherry-RabA]*; Δ*nkuA::argB*, *pyroA4; yA2* | (Qiu et al., 2019) |
| XX222 | GFP-*nudA*<sup>HC</sup>; *argB2::[argB\*-alcAp::mCherry-RabA]*; Δ*nkuA::argB*; *pantoB100; yA2* | (Zhang et al., 2014) |
| XY41 | p25-GFP-Afp<sub>pyrG</sub>; Δ*nkuA::argB*; *pyrG89; pyroA4* | (Zhang et al., 2014) |
| XY42 | *argB2::[argB\*-alcAp::mCherry-RabA]*; Δ*nkuA::argB*; *pyrG89; pantoB100; yA2* | (Qiu et al., 2018) |
| XY136 | Δ*vezA*-Afp<sub>pyrG</sub>; GFP-*nudA*<sup>HC</sup>; *argB2::[argB\*-alcAp::mCherry-RabA]*; *pyrG89; \Delta nkuA::argB; pyroA4; yA2* | (Yao et al., 2015) |
| XY163 | *vezA*-GFP-Afp<sub>pyrG</sub>; *argB2::[argB\*-alcAp::mCherry-RabA]*; *pyrG89; \Delta nkuA::argB pyroA4, wA2* | (Yao et al., 2015) |
| XY167 | ΔTM-*vezA*-GFP-Afp<sub>pyrG</sub>; *argB2::[argB\*-alcA(p)::mCherry-RabA]*; Δ*nkuA::argB*; *pyrG89; pyroA4; wA2* | (Yao et al., 2015) |
| JZ704 | Δ*vezA*-Afp<sub>pyrG</sub>; p150-GFP-Afp<sub>pyrG</sub>; *argB2::[argB\*-alcAp::mCherry-RabA]*; *pyrG89; \Delta nkuA::argB* | This work |
| JZ706 | p150-GFP-Afp<sub>pyrG</sub>; *argB2::[argB\*-alcAp::mCherry-RabA]*; *pyrG89; \Delta nkuA::argB* | This work |
| JZ790 | p62-GFP-Afp<sub>pyrG</sub>, *argB2::[argB\*-alcAp::mCherry-RabA]*; Δ*nkuA::argB*; *pyrG89; pantoB100; yA2* | This work |
| JZ872 | *gpdA*-ΔTM-*vezA*-GFP-Afp<sub>pyrG</sub>; *argB2::[argB\*-alcAp::mCherry-RabA]*; *pyrG89; \Delta nkuA::argB; pantoB100; yA2* | This work |
| JZ942 | Arp11-GFP-Afp<sub>pyrG</sub>; Δ*vezA*-Afp<sub>pyrG</sub>; *argB2::[argB\*-alcAp::mCherry-rabA]*; Δ*nkuA::argB*; *pyrG89; yA2* | This work |
| JZ968 | p150-GFP-Afp<sub>pyrG</sub>; Δ*vezA*-Afp<sub>pyrG</sub>; Δ*hookA*-Afp<sub>pyrG</sub>; *argB2::[argB\*-alcAp::mCherry-RabA]*; Δ*nkuA::argB*; *pyrG89* | This work |
| JZ971 | Arp11-GFP-Afp<sub>pyrG</sub>; Δ*vezA*-Afp<sub>pyrG</sub>; Δ*hookA*-Afp<sub>pyrG</sub>; *argB2::[argB\*-alcAp::mCherry-RabA]*; Δ*nkuA::argB*; *pyrG89* | This work |
| JZ972 | p62-GFP-Afp<sub>pyrG</sub>; Δ*vezA*-Afp<sub>pyrG</sub>; Δ*hookA*-Afp<sub>pyrG</sub>; *argB2::[argB\*-alcAp::mCherry-RabA]*; Δ*nkuA::argB*; *pyrG89; yA2* | This work |
| JZ973 | p25-GFP-Afp<sub>pyrG</sub>; Δ*hookA*-Afp<sub>pyrG</sub>; *argB2::[argB\*-alcAp::mCherry-RabA]*; Δ*nkuA::argB*; *pyrG89; yA2* | This work |
| JZ974 | p25-GFP-Afp<sub>pyrG</sub>; Δ*vezA*-Afp<sub>pyrG</sub>; *argB2::[argB\*-alcAp::mCherry-RabA]*; Δ*nkuA::argB*; *pyrG89* | This work |
| JZ975 | p25-GFP-Afp<sub>pyrG</sub>; Δ*vezA*-Afp<sub>pyrG</sub>; Δ*hookA*-Afp<sub>pyrG</sub>; *argB2::[argB\*-alcAp::mCherry-RabA]*; Δ*nkuA::argB*; *pyrG89* | This work |
| JZ976 | p62-GFP-Afp<sub>pyrG</sub>; Δ*hookA*-Afp<sub>pyrG</sub>; *argB2::[argB\*-alcAp::mCherry-* | This work |
| | *RabA*]; $\Delta nkuA::argB$; *pyrG89*; *wA2* | |
| JZ981 | *p62-GFP-AfpYrG*; $\Delta vezA-AfpYrG$; *argB2::[argB\*-alcAp::mCherry-RabA]*; $\Delta nkuA::argB$; *pyrG89* | This work |
| JZ992 | *p150-GFP-AfpYrG*; *p25-S-AfpYrG*; *argB2::[argB\*-alcA(p)::mCherry-RabA]*; $\Delta nkuA::argB$; *pyrG89* | This work |
| JZ996 | *p150-GFP-AfpYrG*; $\Delta vezA-AfpYrG$; *p25-S-AfpYrG*; $\Delta nkuA::argB$; *pyrG89* | This work |
| JZ1018 | *p25-S-AfpYrG*; *gpdA-ΔTM-vezA-GFP*; *argB2::[argB\*-alcA(p)::mCherry-RabA]*; $\Delta nkuA::argB$; *pyrG89*; *wA2* | This work |
| JZ1025 | *alcA-Arp11*; *argB2::[argB\*-alcA(p)::mCherry-RabA]*; $\Delta nkuA::argB$; *pyrG89*; *pantoB100*; *yA2* | This work |
| JZ1026 | *alcA-Arp1*; *argB2::[argB\*-alcA(p)::mCherry-RabA]*; $\Delta nkuA::argB$; *pyrG89*; *pantoB100*; *yA2* | This work |
| JZ1034 | *alcA-p150*; *argB2::[argB\*-alcA(p)::mCherry-RabA]*; $\Delta nkuA::argB$; *pyrG89*; *pantoB100*; *yA2* | This work |
| JZ1057 | *vezA<sup>Δ563-615</sup>-GFP-AfpYrG*; *argB2::[argB\*-alcAp::mCherry-rabA]*; $\Delta nkuA::argB$; *pyrG89*; *pantoB100*; *yA2* | This work |
| JZ1083 | *gpdA-ΔTM-vezA-GFP*; *alcA-p50*; *argB2::[argB\*-alcA(p)::mCherry-RabA]*; $\Delta nkuA::argB$; *pyrG89*; *yA2* | This work |
| JZ1084 | *gpdA-ΔTM-vezA-GFP*; *alcA-Arp1*; *p25-S-AfpYrG*; *argB2::[argB\*-alcA(p)::mCherry-RabA]*; $\Delta nkuA::argB$; *pyrG89*; *yA2* | This work |
| JZ1085 | *p25-GFP-AfpYrG*; *alcA-p50*; *argB2::[argB\*-alcAp::mCherry-rabA]*; $\Delta nkuA::argB$; *pyrG89*; *yA2* | This work |
| JZ1095 | *p50-GFP-AfpYrG*; $\Delta vezA-AfpYrG$; *argB2::[argB\*-alcAp::mCherry-rabA]*; $\Delta nkuA::argB$; *pyrG89*; *wA2* | This work |
| JZ1108 | *vezA<sup>Δ1-20</sup>-GFP-AfpYrG*; *argB2::[argB\*-alcAp::mCherry-rabA]*; $\Delta nkuA::argB$; *pyrG89*; *pyroA4*; *wA2* | This work |
| JZ1117 | *p25-GFP-AfpYrG*; *p50-S-AfpYrG*; *argB2::[argB\*-alcAp::mCherry-rabA]*; $\Delta nkuA::argB$; *pyrG89* | This work |
| JZ1121 | *p25-GFP-AfpYrG*; *alcA-p150*; *p50-S-AfpYrG*; *argB2::[argB\*-alcAp::mCherry-rabA]*; $\Delta nkuA::argB$; *pyrG89*; *yA2* | This work |
| JZ1122 | *Arp11-GFP-AfpYrG*; *p50-S-AfpYrG*; *argB2::[argB\*-alcAp::mCherry-rabA]*; $\Delta nkuA::argB$; *pyrG89* | This work |
| JZ1123 | *Arp11-GFP-AfpYrG*; *p50-S-AfpYrG*; $\Delta vezA-AfpYrG$; *argB2::[argB\*-alcAp::mCherry-rabA]*; $\Delta nkuA::argB$; *pyrG89* | This work |
| JZ1125 | *p50-GFP-AfpYrG*; *alcA-Arp1*; *argB2::[argB\*-alcAp::mCherry-rabA]*; $\Delta nkuA::argB$; *pyrG89*; *yA2* | This work |
| RQ70 | *p25-S-AfpYrG*; *argB2::[argB\*-alcAp::mCherry-rabA]*; $\Delta nkuA::argB$; *pyrG89*; *pyroA4*; *wA2* | This work |
| SX1 | *p50-S-AfpYrG*; *argB2::[argB\*-alcAp::mCherry-rabA]*; $\Delta nkuA::argB$; *pyrG89*; *pantoB100*; *yA2* | This work |
| SX2 | *p50-GFP-AfpYrG*; *argB2::[argB\*-alcAp::mCherry-rabA]*; $\Delta nkuA::argB$; *pyrG89*; *pyroA4*; *wA2* | This work |
| XX365 | $\Delta vezA-AfpYrG$; *GFP-histone H1 (hhoA-GFP-Afribo)*; *argB2::[argB\*-alcAp::mCherry-RabA]*; possibly *pyrG89*; possibly $\Delta nkuA::argB$; *wA2* | This work |
| XX367 | *GFP-histone H1 (hhoA-GFP-Afribo)*; *argB2::[argB\*-alcAp::mCherry-RabA]*; possibly *pyrG89*; possibly $\Delta nkuA::argB$ | This work |
| XX370 | *p150-GFP-AfpYrG*; *argB2::[argB\*-alcAp::mCherry-RabA]* | This work |
| XX371 | $\Delta vezA-AfpYrG$; *p150-GFP-AfpYrG*; *argB2::[argB\*-alcAp::mCherry-* | This work |
|  | *RabA]* |  |
| XX374 | ΔvezA-Afp<sub>pyr</sub>G; *argB2::[argB<sup>-</sup>-alcAp::mCherry-RabA]*; ΔnkuA::argB, *pyrG89; pantoB100* | This work |
| XX518 | ΔvezA-Afp<sub>pyr</sub>G; GFP-*nudA*<sup>HC</sup>; *gpdA-ΔC-hookA-S-Afp<sub>pyr</sub>G; pyrG89; yA1, ΔnkuA::argB* | This work |
| XX618 | GFP-*nudA*<sup>R1602E, K1645E</sup>; ΔvezA-Afp<sub>pyr</sub>G; *argB2::[argB<sup>-</sup>-alcAp::mCherry-RabA]*; ΔnkuA::argB, *pyrG89;(RQ294x JZ698)* | This work |
| XX721 | p150-GFP-Afp<sub>pyr</sub>G; ΔhookA-Afp<sub>pyr</sub>G; *argB2::[argB<sup>-</sup>-alcAp::mCherry-RabA]* | This work |
| XX723 | p150-GFP-Afp<sub>pyr</sub>G; ΔhookA-Afp<sub>pyr</sub>G; ΔvezA-Afp<sub>pyr</sub>G; *argB2::[argB<sup>-</sup>-alcAp::mCherry-RabA]* | This work |
| XX760 | Arp11-GFP-Afp<sub>pyr</sub>G; ΔhookA-Afp<sub>pyr</sub>G; *argB2::[argB<sup>-</sup>-alcAp::mCherry-RabA]* | This work |
| XX762 | Arp11-GFP-Afp<sub>pyr</sub>G; ΔhookA-Afp<sub>pyr</sub>G; ΔvezA-Afp<sub>pyr</sub>G; *argB2::[argB<sup>-</sup>-alcAp::mCherry-RabA]* | This work |
| XX1017 | p50-GFP-afp<sub>pyr</sub>G; ΔhookA-Afp<sub>pyr</sub>G; *argB2::[argB<sup>-</sup>-alcAp::mCherry-RabA]*; wA2 | This work |
| XX1018 | p50-GFP-afp<sub>pyr</sub>G; ΔhookA-Afp<sub>pyr</sub>G; ΔvezA-Afp<sub>pyr</sub>G; *argB2::[argB<sup>-</sup>-alcAp::mCherry-RabA]*; wA2 | This work |

## References

Abenza, J.F., A. Pantazopoulou, J.M. Rodriguez, A. Galindo, and M.A. Penalva. 2009. Long-distance movement of Aspergillus nidulans early endosomes on microtubule tracks. Traffic. 10:57–75.

Amaro, I.A., M. Costanzo, C. Boone, and T.C. Huffaker. 2008. The Saccharomyces cerevisiae homolog of p24 is essential for maintaining the association of p150Glued with the dynactin complex. Genetics. 178:703–709.

Ananthanarayanan, V., M. Schattat, S.K. Vogel, A. Krull, N. Pavin, and I.M. Tolić-Nørrelykke. 2013. Dynein motion switches from diffusive to directed upon cortical anchoring. Cell. 153:1526–1536.

Bahloul, A., M.C. Simmler, V. Michel, M. Leibovici, I. Perfettini, I. Roux, D. Weil, S. Nouaille, J. Zuo, C. Zadro, D. Licastro, P. Gasparini, P. Avan, J.P. Hardelin, and C. Petit. 2009. Vezatin, an integral membrane protein of adherens junctions, is required for the sound resilience of cochlear hair cells. EMBO Mol Med. 1:125–138.

Baumann, S., J. König, J. Koepke, and M. Feldbrügge. 2014. Endosomal transport of septin mRNA and protein indicates local translation on endosomes and is required for correct septin filamentation. EMBO Rep. 15:94–102.

Baumbach, J., A. Murthy, M.A. McClintock, C.I. Dix, R. Zalyte, H.T. Hoang, and S.L. Bullock. 2017. Lissencephaly-1 is a context-dependent regulator of the human dynein complex. Elife. 6.

Bieger, B.D., A.H. Osmani, X. Xiang, and M.J. Egan. 2021. The spindle pole-body localization of activated cytoplasmic dynein is cell cycle-dependent in Aspergillus nidulans. Fungal Genet Biol. 148:103519.

Bielska, E., M. Schuster, Y. Roger, A. Berepiki, D.M. Soanes, N.J. Talbot, and G. Steinberg. 2014. Hook is an adapter that coordinates kinesin-3 and dynein cargo attachment on early endosomes. J Cell Biol. 204:989–1007.

Bingham, J.B., and T.A. Schroer. 1999. Self-regulated polymerization of the actin-related protein Arp1. Curr Biol. 9:223–226.

Callejas-Negrete, O.A., M. Plamann, R. Schnittker, S. Bartnicki-García, R.W. Roberson, G. Pimienta, and R.R. Mouriño-Pérez. 2015. Two microtubule-plus-end binding proteins LIS1-1 and LIS1-2, homologues of human LIS1 in Neurospora crassa. Fungal Genet Biol. 82:213–227.

Chaaban, S., and A.P. Carter. 2022. Structure of dynein-dynactin on microtubules shows tandem adaptor binding. Nature. 610:212–216.

Cheong, F.K., L. Feng, A. Sarkeshik, J.R. Yates, 3rd, and T.A. Schroer. 2014. Dynactin integrity depends upon direct binding of dynamitin to Arp1. Mol Biol Cell. 25:2171-2180.

Chowdhury, S., S.A. Ketcham, T.A. Schroer, and G.C. Lander. 2015. Structural organization of the dynein-dynactin complex bound to microtubules. Nat Struct Mol Biol. 22:345–347.

Christensen, J.R., A.A. Kendrick, J.B. Truong, A. Aguilar-Maldonado, V. Adani, M. Dzieciatkowska, and S.L. Reck-Peterson. 2021. Cytoplasmic dynein-1 cargo diversity is mediated by the combinatorial assembly of FTS-Hook-FHIP complexes. Elife. 10.

Clark, S.W., and M.D. Rose. 2006. Arp10p is a pointed-end-associated component of yeast dynactin. Mol Biol Cell. 17:738–748.

Danglot, L., T. Freret, N. Le Roux, N. Narboux Neme, A. Burgo, V. Hyenne, A. Roumier, V. Contremoulins, F. Dauphin, J.C. Bizot, G. Vodjdani, P. Gaspar, M. Boulouard, J.C. Poncer, T. Galli, and M.C. Simmler. 2012. Vezatin is essential for dendritic spine morphogenesis and functional synaptic maturation. J Neurosci. 32:9007–9022.

De Pace, R., S. Ghosh, V.H. Ryan, M. Sohn, M. Jarnik, P. Rezvan Sangsari, N.Y. Morgan, R.K. Dale, M.E. Ward, and J.S. Bonifacino. 2024. Messenger RNA transport on lysosomal vesicles maintains axonal mitochondrial homeostasis and prevents axonal degeneration. Nat Neurosci.

Drerup, C.M., A.L. Herbert, K.R. Monk, and A.V. Nechiporuk. 2017. Regulation of mitochondria-dynactin interaction and mitochondrial retrograde transport in axons. Elife. 6.

Echeverri, C.J., B.M. Paschal, K.T. Vaughan, and R.B. Vallee. 1996. Molecular characterization of the 50-kD subunit of dynactin reveals function for the complex in chromosome alignment and spindle organization during mitosis. J Cell Biol. 132:617–633.

Eckley, D.M., S.R. Gill, K.A. Melkonian, J.B. Bingham, H.V. Goodson, J.E. Heuser, and T.A. Schroer. 1999. Analysis of dynactin subcomplexes reveals a novel actin-related protein associated with the arp1 minifilament pointed end. J Cell Biol. 147:307–320.

Egan, M.J., M.A. McClintock, I.H. Hollyer, H.L. Elliott, and S.L. Reck-Peterson. 2015. Cytoplasmic dynein is required for the spatial organization of protein aggregates in filamentous fungi. Cell Rep. 11:201-209.

Egan, M.J., K. Tan, and S.L. Reck-Peterson. 2012. Lis1 is an initiation factor for dynein-driven organelle transport. J Cell Biol. 197:971–982.

Elshenawy, M.M., E. Kusakci, S. Volz, J. Baumbach, S.L. Bullock, and A. Yildiz. 2020. Lis1 activates dynein motility by modulating its pairing with dynactin. Nat Cell Biol. 22:570–578.

Eshel, D., L.A. Urrestarazu, S. Vissers, J.C. Jauniaux, J.C. van Vliet-Reedijk, R.J. Planta, and I.R. Gibbons. 1993. Cytoplasmic dynein is required for normal nuclear segregation in yeast. Proc Natl Acad Sci U S A. 90:11172–11176.

Etxebeste, O., M. Villarino, A. Markina-Iñarrairaegui, L. Araújo-Bazán, and E.A. Espeso. 2013. Cytoplasmic dynamics of the general nuclear import machinery in apically growing syncytial cells. PLoS One. 8:e85076.

Fagarasanu, A., M. Fagarasanu, G.A. Eitzen, J.D. Aitchison, and R.A. Rachubinski. 2006. The peroxisomal membrane protein Inp2p is the peroxisome-specific receptor for the myosin V motor Myo2p of Saccharomyces cerevisiae. Dev Cell. 10:587–600.

Gama, J.B., C. Pereira, P.A. Simões, R. Celestino, R.M. Reis, D.J. Barbosa, H.R. Pires, C. Carvalho, J. Amorim, A.X. Carvalho, D.K. Cheerambathur, and R. Gassmann. 2017. Molecular mechanism of dynein recruitment to kinetochores by the Rod-Zw10-Zwilch complex and Spindly. J Cell Biol. 216:943–960.

Garces, J.A., I.B. Clark, D.I. Meyer, and R.B. Vallee. 1999. Interaction of the p62 subunit of dynactin with Arp1 and the cortical actin cytoskeleton. Curr Biol. 9:1497–1500.

Gill, S.R., T.A. Schroer, I. Szilak, E.R. Steuer, M.P. Sheetz, and D.W. Cleveland. 1991. Dynactin, a conserved, ubiquitously expressed component of an activator of vesicle motility mediated by cytoplasmic dynein. J Cell Biol. 115:1639–1650.

Go, C.D., J.D.R. Knight, A. Rajasekharan, B. Rathod, G.G. Hesketh, K.T. Abe, J.Y. Youn, P. Samavarchi-Tehrani, H. Zhang, L.Y. Zhu, E. Popiel, J.P. Lambert, É. Coyaud, S.W.T. Cheung, D. Rajendran, C.J. Wong, H. Antonicka, L. Pelletier, A.F. Palazzo, E.A. Shoubridge, B. Raught, and A.C. Gingras. 2021. A proximity-dependent biotinylation map of a human cell. Nature. 595:120–124.

Grotjahn, D.A., S. Chowdhury, Y. Xu, R.J. McKenney, T.A. Schroer, and G.C. Lander. 2018. Cryo-electron tomography reveals that dynactin recruits a team of dyneins for processive motility. Nat Struct Mol Biol. 25:203–207.

Guimaraes, S.C., M. Schuster, E. Bielska, G. Dagdas, S. Kilaru, B.R. Meadows, M. Schrader, and G. Steinberg. 2015. Peroxisomes, lipid droplets, and endoplasmic reticulum "hitchhike" on motile early endosomes. J Cell Biol. 211:945–954.

Guo, X., G.G. Farias, R. Mattera, and J.S. Bonifacino. 2016. Rab5 and its effector FHF contribute to neuronal polarity through dynein-dependent retrieval of somatodendritic proteins from the axon. Proc Natl Acad Sci U S A. 113:E5318–5327.

Haghnia, M., V. Cavalli, S.B. Shah, K. Schimmelpfeng, R. Brusch, G. Yang, C. Herrera, A. Pilling, and L.S. Goldstein. 2007. Dynactin is required for coordinated bidirectional motility, but not for dynein membrane attachment. Mol Biol Cell. 18:2081–2089.

Han, G., B. Liu, J. Zhang, W. Zuo, N.R. Morris, and X. Xiang. 2001. The Aspergillus cytoplasmic dynein heavy chain and NUDF localize to microtubule ends and affect microtubule dynamics. Curr Biol. 11:719–724.

Hein, M.Y., N.C. Hubner, I. Poser, J. Cox, N. Nagaraj, Y. Toyoda, I.A. Gak, I. Weisswange, J. Mansfeld, F. Buchholz, A.A. Hyman, and M. Mann. 2015. A human interactome in three quantitative dimensions organized by stoichiometries and abundances. Cell. 163:712–723.

Higuchi, Y., P. Ashwin, Y. Roger, and G. Steinberg. 2014. Early endosome motility spatially organizes polysome distribution. J Cell Biol. 204:343–357.

Holdsworth-Carson, S.J., J.N. Fung, H.T. Luong, Y. Sapkota, L.M. Bowdler, L. Wallace, W.T. Teh, J.E. Powell, J.E. Girling, M. Healey, G.W. Montgomery, and P.A. Rogers. 2016. Endometrial vezatin and its association with endometriosis risk. Hum Reprod. 31:999–1013.

Holzbaur, E.L., J.A. Hammarback, B.M. Paschal, N.G. Kravit, K.K. Pfister, and R.B. Vallee. 1991. Homology of a 150K cytoplasmic dynein-associated polypeptide with the Drosophila gene Glued. Nature. 351:579–583.

Htet, Z.M., J.P. Gillies, R.W. Baker, A.E. Leschziner, M.E. DeSantis, and S.L. Reck-Peterson. 2020. LIS1 promotes the formation of activated cytoplasmic dynein-1 complexes. Nat Cell Biol. 22:518–525.

Jha, R., J. Roostalu, N.I. Cade, M. Trokter, and T. Surrey. 2017. Combinatorial regulation of the balance between dynein microtubule end accumulation and initiation of directed motility. Embo j. 36:3387–3404.

Karasmanis, E.P., J.M. Reimer, A.A. Kendrick, K.H.V. Nguyen, J.A. Rodriguez, J.B. Truong, I. Lahiri, S.L. Reck-Peterson, and A.E. Leschziner. 2023. Lis1 relieves cytoplasmic dynein-1 autoinhibition by acting as a molecular wedge. Nat Struct Mol Biol. 30:1357–1364.

Karki, S., and E.L. Holzbaur. 1995. Affinity chromatography demonstrates a direct binding between cytoplasmic dynein and the dynactin complex. J Biol Chem. 270:28806–28811.

Karki, S., M.K. Tokito, and E.L. Holzbaur. 2000. A dynactin subunit with a highly conserved cysteine-rich motif interacts directly with Arp1. J Biol Chem. 275:4834–4839.

Kim, H., S.C. Ling, G.C. Rogers, C. Kural, P.R. Selvin, S.L. Rogers, and V.I. Gelfand. 2007. Microtubule binding by dynactin is required for microtubule organization but not cargo transport. J Cell Biol. 176:641–651.

King, S.J., C.L. Brown, K.C. Maier, N.J. Quintyne, and T.A. Schroer. 2003. Analysis of the dynein-dynactin interaction in vitro and in vivo. Mol Biol Cell. 14:5089–5097.

Koppel, N., M.B. Friese, H.L. Cardasis, T.A. Neubert, and S.J. Burden. 2019. Vezatin is required for the maturation of the neuromuscular synapse. Mol Biol Cell. 30:2571–2583.

Kussel-Andermann, P., A. El-Amraoui, S. Safieddine, S. Nouaille, I. Perfettini, M. Lecuit, P. Cossart, U. Wolfrum, and C. Petit. 2000. Vezatin, a novel transmembrane protein, bridges myosin VIIA to the cadherin-catenins complex. EMBO J. 19:6020–6029.

Lammers, L.G., and S.M. Markus. 2015. The dynein cortical anchor Num1 activates dynein motility by relieving Pac1/LIS1-mediated inhibition. J Cell Biol. 211:309–322.

Lau, C.K., F.J. O’Reilly, B. Santhanam, S.E. Lacey, J. Rappsilber, and A.P. Carter. 2021. Cryo-EM reveals the complex architecture of dynactin’s shoulder region and pointed end. Embo j. 40:e106164.

Lee, I.H., S. Kumar, and M. Plamann. 2001. Null mutants of the neurospora actin-related protein 1 pointed-end complex show distinct phenotypes. Mol Biol Cell. 12:2195–2206.

Lee, W.L., J.R. Oberle, and J.A. Cooper. 2003. The role of the lissencephaly protein Pac1 during nuclear migration in budding yeast. J Cell Biol. 160:355–364.

Lenz, J.H., I. Schuchardt, A. Straube, and G. Steinberg. 2006. A dynein loading zone for retrograde endosome motility at microtubule plus-ends. Embo j. 25:2275–2286.

Li, Y.S., Y.Z. Chen, X.B. Guo, X. Liu, and L.P. Li. 2015. VEZT as a novel independent prognostic factor in gastric cancer. Cancer Biomark. 15:375–380.

Li, Y.Y., E. Yeh, T. Hays, and K. Bloom. 1993. Disruption of mitotic spindle orientation in a yeast dynein mutant. Proc Natl Acad Sci U S A. 90:10096–10100.

Lloyd, T.E., J. Machamer, K. O’Hara, J.H. Kim, S.E. Collins, M.Y. Wong, B. Sahin, W. Imlach, Y. Yang, E.S. Levitan, B.D. McCabe, and A.L. Kolodkin. 2012. The p150(Glued) CAP-Gly domain regulates initiation of retrograde transport at synaptic termini. Neuron. 74:344–360.

Maier, K.C., J.E. Godfrey, C.J. Echeverri, F.K. Cheong, and T.A. Schroer. 2008. Dynamitin mutagenesis reveals protein-protein interactions important for dynactin structure. Traffic. 9:481–491.

Markus, S.M., M.G. Marzo, and R.J. McKenney. 2020. New insights into the mechanism of dynein motor regulation by lissencephaly-1. Elife. 9.

Marzo, M.G., J.M. Griswold, and S.M. Markus. 2020. Pac1/LIS1 stabilizes an uninhibited conformation of dynein to coordinate its localization and activity. Nat Cell Biol. 22:559–569.

McKenney, R.J., W. Huynh, M.E. Tanenbaum, G. Bhabha, and R.D. Vale. 2014. Activation of cytoplasmic dynein motility by dynactin-cargo adapter complexes. Science. 345:337–341.

Minke, P.F., I.H. Lee, J.H. Tinsley, K.S. Bruno, and M. Plamann. 1999. Neurospora crassa ro-10 and ro-11 genes encode novel proteins required for nuclear distribution. Mol Microbiol. 32:1065–1076.

Mirdita, M., K. Schütze, Y. Moriwaki, L. Heo, S. Ovchinnikov, and M. Steinegger. 2022. ColabFold: making protein folding accessible to all. Nat Methods. 19:679–682.

Moore, J.K., J. Li, and J.A. Cooper. 2008. Dynactin function in mitotic spindle positioning. Traffic. 9:510–527.

Morita, Y., K. Takegawa, B.M. Collins, and Y. Higuchi. 2024. Polarity-dependent expression and localization of secretory glucoamylase mRNA in filamentous fungal cells. Microbiol Res. 282:127653.

Moughamian, A.J., and E.L. Holzbaur. 2012. Dynactin is required for transport initiation from the distal axon. Neuron. 74:331–343.

Nayak, T., E. Szewczyk, C.E. Oakley, A. Osmani, L. Ukil, S.L. Murray, M.J. Hynes, S.A. Osmani, and B.R. Oakley. 2006. A versatile and efficient gene-targeting system for Aspergillus nidulans. Genetics. 172:1557–1566.

Oakley, B.R., C.E. Oakley, Y. Yoon, and M.K. Jung. 1990. Gamma-tubulin is a component of the spindle pole body that is essential for microtubule function in Aspergillus nidulans. Cell. 61:1289–1301.

Okada, K., B.R. Iyer, L.G. Lammers, P.A. Gutierrez, W. Li, S.M. Markus, and R.J. McKenney. 2023. Conserved roles for the dynein intermediate chain and Ndel1 in assembly and activation of dynein. Nat Commun. 14:5833.

Olenick, M.A., R. Dominguez, and E.L.F. Holzbaur. 2019. Dynein activator Hook1 is required for trafficking of BDNF-signaling endosomes in neurons. J Cell Biol. 218:220–233.

Otamendi, A., E. Perez-de-Nanclares-Arregi, E. Oiartzabal-Arano, M.S. Cortese, E.A. Espeso, and O. Etxebeste. 2019. Developmental regulators FlbE/D orchestrate the polarity site-to-nucleus dynamics of the fungal bZIP transcription factor FlbB. Cell Mol Life Sci. 76:4369–4390.

Pagliardini, L., D. Gentilini, A.M. Sanchez, M. Candiani, P. Viganò, and A.M. Di Blasio. 2015. Replication and meta-analysis of previous genome-wide association studies confirm vezatin as the locus with the strongest evidence for association with endometriosis. Hum Reprod. 30:987–993.

Penalva, M.A., J. Zhang, X. Xiang, and A. Pantazopoulou. 2017. Transport of fungal RAB11 secretory vesicles involves myosin-5, dynein/dynactin/p25, and kinesin-1 and is independent of kinesin-3. Mol Biol Cell. 28:947–961.

Plamann, M., P.F. Minke, J.H. Tinsley, and K.S. Bruno. 1994. Cytoplasmic dynein and actin-related protein Arp1 are required for normal nuclear distribution in filamentous fungi. J Cell Biol. 127:139–149.

Pohlmann, T., S. Baumann, C. Haag, M. Albrecht, and M. Feldbrugge. 2015. A FYVE zinc finger domain protein specifically links mRNA transport to endosome trafficking. Elife. 4.

Qiu, R., J. Zhang, and X. Xiang. 2013. Identification of a novel site in the tail of dynein heavy chain important for dynein function in vivo. J Biol Chem. 288:2271–2280.

Qiu, R., J. Zhang, and X. Xiang. 2018. p25 of the dynactin complex plays a dual role in cargo binding and dynactin regulation. J Biol Chem. 293:15606–15619.

Qiu, R., J. Zhang, and X. Xiang. 2019. LIS1 regulates cargo-adapter-mediated activation of dynein by overcoming its autoinhibition in vivo. J Cell Biol. 218:3630–3646.

Qiu, R., J. Zhang, and X. Xiang. 2020. The splicing-factor Prp40 affects dynein-dynactin function in Aspergillus nidulans. Mol Biol Cell. 31:1289–1301.

Rao, L., and A. Gennerich. 2024. Structure and Function of Dynein’s Non-Catalytic Subunits. Cells. 13.

Saito, K., T. Murayama, T. Hata, T. Kobayashi, K. Shibata, S. Kazuno, T. Fujimura, T. Sakurai, and Y.Y. Toyoshima. 2020. Conformational diversity of dynactin sidearm and domain organization of its subunit p150. Mol Biol Cell. 31:1218–1231.

Salogiannis, J., J.R. Christensen, L.D. Songster, A. Aguilar-Maldonado, N. Shukla, and S.L. Reck-Peterson. 2021. PxdA interacts with the DipA phosphatase to regulate peroxisome hitchhiking on early endosomes. Mol Biol Cell. 32:492–503.

Salogiannis, J., M.J. Egan, and S.L. Reck-Peterson. 2016. Peroxisomes move by hitchhiking on early endosomes using the novel linker protein PxdA. J Cell Biol. 212:289–296.

Sanda, M., N. Ohara, A. Kamata, Y. Hara, H. Tamaki, J. Sukegawa, T. Yanagisawa, K. Fukunaga, H. Kondo, and H. Sakagami. 2010. Vezatin, a potential target for ADP-ribosylation factor 6, regulates the dendritic formation of hippocampal neurons. Neurosci Res. 67:126–136.

Schafer, D.A., S.R. Gill, J.A. Cooper, J.E. Heuser, and T.A. Schroer. 1994. Ultrastructural analysis of the dynactin complex: an actin-related protein is a component of a filament that resembles F-actin. J Cell Biol. 126:403–412.

Schlager, M.A., H.T. Hoang, L. Urnavicius, S.L. Bullock, and A.P. Carter. 2014. In vitro reconstitution of a highly processive recombinant human dynein complex. EMBO J. 33:1855–1868.

Schroeder, C.M., and R.D. Vale. 2016. Assembly and activation of dynein-dynactin by the cargo adaptor protein Hook3. J Cell Biol. 214:309–318.

Schroer, T.A. 2004. Dynactin. Annu Rev Cell Dev Biol. 20:759–779.

Seiler, S., M. Plamann, and M. Schliwa. 1999. Kinesin and dynein mutants provide novel insights into the roles of vesicle traffic during cell morphogenesis in Neurospora. Curr Biol. 9:779–785.

Sheeman, B., P. Carvalho, I. Sagot, J. Geiser, D. Kho, M.A. Hoyt, and D. Pellman. 2003. Determinants of S. cerevisiae dynein localization and activation: implications for the mechanism of spindle positioning. Curr Biol. 13:364–372.

Singh, K., C.K. Lau, G. Manigrasso, J.B. Gama, R. Gassmann, and A.P. Carter. 2024. Molecular mechanism of dynein-dynactin complex assembly by LIS1. Science. 383:eadk8544.

Songster, L.D., D. Bhuyan, J.R. Christensen, and S.L. Reck-Peterson. 2023. Woronin body hitchhiking on early endosomes is dispensable for septal localization in Aspergillus nidulans. Mol Biol Cell. 34:br9.

Sousa, S., D. Cabanes, A. El-Amraoui, C. Petit, M. Lecuit, and P. Cossart. 2004. Unconventional myosin VIIa and vezatin, two proteins crucial for Listeria entry into epithelial cells. J Cell Sci. 117:2121–2130.

Spinner, M.A., K. Pinter, C.M. Drerup, and T.G. Herman. 2020. A Conserved Role for Vezatin Proteins in Cargo-Specific Regulation of Retrograde Axonal Transport. Genetics. 216:431–445.

Splinter, D., D.S. Razafsky, M.A. Schlager, A. Serra-Marques, I. Grigoriev, J. Demmers, N. Keijzer, K. Jiang, I. Poser, A.A. Hyman, C.C. Hoogenraad, S.J. King, and A. Akhmanova. 2012. BICD2, dynactin, and LIS1 cooperate in regulating dynein recruitment to cellular structures. Mol Biol Cell. 23:4226–4241.

Szewczyk, E., T. Nayak, C.E. Oakley, H. Edgerton, Y. Xiong, N. Taheri-Talesh, S.A. Osmani, and B.R. Oakley. 2006. Fusion PCR and gene targeting in Aspergillus nidulans. Nat Protoc. 1:3111–3120.

Torisawa, T., M. Ichikawa, A. Furuta, K. Saito, K. Oiwa, H. Kojima, Y.Y. Toyoshima, and K. Furuta. 2014. Autoinhibition and cooperative activation mechanisms of cytoplasmic dynein. Nat Cell Biol. 16:1118–1124.

Tripathy, S.K., S.J. Weil, C. Chen, P. Anand, R.B. Vallee, and S.P. Gross. 2014. Autoregulatory mechanism for dynactin control of processive and diffusive dynein transport. Nat Cell Biol. 16:1192–1201.

Twelvetrees, A.E., S. Pernigo, A. Sanger, P. Guedes-Dias, G. Schiavo, R.A. Steiner, M.P. Dodding, and E.L. Holzbaur. 2016. The Dynamic Localization of Cytoplasmic Dynein in Neurons Is Driven by Kinesin-1. Neuron. 90:1000–1015.

Urnavicius, L., C.K. Lau, M.M. Elshenawy, E. Morales-Rios, C. Motz, A. Yildiz, and A.P. Carter. 2018. Cryo-EM shows how dynactin recruits two dyneins for faster movement. Nature. 554:202–206.

Urnavicius, L., K. Zhang, A.G. Diamant, C. Motz, M.A. Schlager, M. Yu, N.A. Patel, C.V. Robinson, and A.P. Carter. 2015. The structure of the dynactin complex and its interaction with dynein. Science. 347:1441–1446.

Valetti, C., D.M. Wetzel, M. Schrader, M.J. Hasbani, S.R. Gill, T.E. Kreis, and T.A. Schroer. 1999. Role of dynactin in endocytic traffic: effects of dynamitin overexpression and colocalization with CLIP-170. Mol Biol Cell. 10:4107–4120.

Vaughan, K.T., S.H. Tynan, N.E. Faulkner, C.J. Echeverri, and R.B. Vallee. 1999. Colocalization of cytoplasmic dynein with dynactin and CLIP-170 at microtubule distal ends. J Cell Sci. 112 (Pt 10):1437–1447.

Vaughan, K.T., and R.B. Vallee. 1995. Cytoplasmic dynein binds dynactin through a direct interaction between the intermediate chains and p150Glued. J Cell Biol. 131:1507–1516.

Vaughan, P.S., P. Miura, M. Henderson, B. Byrne, and K.T. Vaughan. 2002. A role for regulated binding of p150(Glued) to microtubule plus ends in organelle transport. J Cell Biol. 158:305–319.

Wang, Y., J. Yuan, X. Yu, X. Liu, C. Tan, Y. Chen, and T. Xu. 2021. Vezatin regulates seizures by controlling AMPAR-mediated synaptic activity. Cell Death Dis. 12:936.

Waring, R.B., G.S. May, and N.R. Morris. 1989. Characterization of an inducible expression system in Aspergillus nidulans using alcA and tubulin-coding genes. Gene. 79:119–130.

Waterman-Storer, C.M., S. Karki, and E.L. Holzbaur. 1995. The p150Glued component of the dynactin complex binds to both microtubules and the actin-related protein centractin (Arp-1). Proc Natl Acad Sci U S A. 92:1634–1638.

Xiang, X., S.M. Beckwith, and N.R. Morris. 1994. Cytoplasmic dynein is involved in nuclear migration in Aspergillus nidulans. Proc Natl Acad Sci U S A. 91:2100–2104.

Xiang, X., G. Han, D.A. Winkelmann, W. Zuo, and N.R. Morris. 2000. Dynamics of cytoplasmic dynein in living cells and the effect of a mutation in the dynactin complex actin-related protein Arp1. Curr Biol. 10:603–606.

Xiang, X., and R. Qiu. 2020. Cargo-Mediated Activation of Cytoplasmic Dynein in vivo. Front Cell Dev Biol. 8:598952.

Xu, L., M.E. Sowa, J. Chen, X. Li, S.P. Gygi, and J.W. Harper. 2008. An FTS/Hook/p107(FHIP) complex interacts with and promotes endosomal clustering by the homotypic vacuolar protein sorting complex. Mol Biol Cell. 19:5059–5071.

Yamada, M., S. Toba, T. Takitoh, Y. Yoshida, D. Mori, T. Nakamura, A.H. Iwane, T. Yanagida, H. Imai, L.Y. Yu-Lee, T. Schroer, A. Wynshaw-Boris, and S. Hirotsune. 2010. mNUDC is required for plus-end-directed transport of cytoplasmic dynein and dynactins by kinesin-1. Embo j. 29:517–531.

Yang, L., L. Ukil, A. Osmani, F. Nahm, J. Davies, C.P. De Souza, X. Dou, A. Perez-Balaguer, and S.A. Osmani. 2004. Rapid production of gene replacement constructs and generation of a green fluorescent protein-tagged centromeric marker in Aspergillus nidulans. Eukaryot Cell. 3:1359–1362.

Yao, X., H.N. Arst, Jr., X. Wang, and X. Xiang. 2015. Discovery of a vezatin-like protein for dynein-mediated early endosome transport. Mol Biol Cell. 26:3816–3827.

Yao, X., X. Wang, and X. Xiang. 2014. FHIP and FTS proteins are critical for dynein-mediated transport of early endosomes in Aspergillus. Mol Biol Cell. 25:2181–2189.

Yao, X., J. Zhang, H. Zhou, E. Wang, and X. Xiang. 2012. In vivo roles of the basic domain of dynactin p150 in microtubule plus-end tracking and dynein function. Traffic. 13:375–387.

Yeh, T.Y., A.K. Kowalska, B.R. Scipioni, F.K. Cheong, M. Zheng, U. Derewenda, Z.S. Derewenda, and T.A. Schroer. 2013. Dynactin helps target Polo-like kinase 1 to kinetochores via its left-handed beta-helical p27 subunit. Embo j. 32:1023–1035.

Yeh, T.Y., N.J. Quintyne, B.R. Scipioni, D.M. Eckley, and T.A. Schroer. 2012. Dynactin’s pointed-end complex is a cargo-targeting module. Mol Biol Cell. 23:3827–3837.

Zander, S., S. Baumann, S. Weidtkamp-Peters, and M. Feldbrügge. 2016. Endosomal assembly and transport of heteromeric septin complexes promote septin cytoskeleton formation. J Cell Sci. 129:2778–2792.

Zhang, J., S. Li, R. Fischer, and X. Xiang. 2003. Accumulation of cytoplasmic dynein and dynactin at microtubule plus ends in Aspergillus nidulans is kinesin dependent. Mol Biol Cell. 14:1479–1488.

Zhang, J., R. Qiu, H.N. Arst, Jr., M.A. Penalva, and X. Xiang. 2014. HookA is a novel dynein-early endosome linker critical for cargo movement in vivo. J Cell Biol. 204:1009–1026.

Zhang, J., R. Qiu, B.D. Bieger, C.E. Oakley, B.R. Oakley, M.J. Egan, and X. Xiang. 2023. Aspergillus SUMOylation mutants exhibit chromosome segregation defects including chromatin bridges. Genetics. 225.

Zhang, J., R. Qiu, and X. Xiang. 2018. The actin capping protein in Aspergillus nidulans enhances dynein function without significantly affecting Arp1 filament assembly. Sci Rep. 8:11419.

Zhang, J., L. Wang, L. Zhuang, L. Huo, S. Musa, S. Li, and X. Xiang. 2008. Arp11 affects dynein-dynactin interaction and is essential for dynein function in Aspergillus nidulans. Traffic. 9:1073–1087.

Zhang, J., X. Yao, L. Fischer, J.F. Abenza, M.A. Penalva, and X. Xiang. 2011. The p25 subunit of the dynactin complex is required for dynein-early endosome interaction. J Cell Biol. 193:1245–1255.

Zhang, J., L. Zhuang, Y. Lee, J.F. Abenza, M.A. Peñalva, and X. Xiang. 2010. The microtubule plus-end localization of Aspergillus dynein is important for dynein-early-endosome interaction but not for dynein ATPase activation. J Cell Sci. 123:3596–3604.

Zhang, K., H.E. Foster, A. Rondelet, S.E. Lacey, N. Bahi-Buisson, A.W. Bird, and A.P. Carter. 2017a. Cryo-EM Reveals How Human Cytoplasmic Dynein Is Auto-inhibited and Activated. Cell. 169:1303–1314.e1318.

Zhang, Y., X. Gao, R. Manck, M. Schmid, A.H. Osmani, S.A. Osmani, N. Takeshita, and R. Fischer. 2017b. Microtubule-organizing centers of Aspergillus nidulans are anchored at septa by a disordered protein. Mol Microbiol. 106:285–303.

Zhao, Y., S. Oten, and A. Yildiz. 2023. Nde1 promotes Lis1-mediated activation of dynein. Nat Commun. 14:7221.

